# Inhibition of proline-rich-tyrosine kinase 2 restores cardioprotection by remote ischemic preconditioning in type 2 diabetes mellitus

**DOI:** 10.1101/2023.04.25.538211

**Authors:** Ralf Erkens, Dragos Duse, Amanda Brum, Alexandra Chadt, Stefanie Becher, Mauro Siragusa, Christine Quast, Johanna Müssig, Michael Roden, Miriam Cortese-Krott, Eckhard Lammert E, Ingrid Fleming, Christian Jung, Hadi Al-Hasani, Gerd Heusch, Malte Kelm

## Abstract

**Background:** Endothelial function and cardioprotection through remote ischemic preconditioning (rIPC) are severely impaired in type 2 diabetes mellitus (T2DM). Proline-rich tyrosine kinase 2 (Pyk2), a downstream target of the insulin receptor, reduces endothelial nitric oxide synthase (eNOS) activity. Therapeutic options to rescue cardioprotection in T2DM and improve outcomes after acute myocardial infarction (AMI) are lacking. We hypothesized that vascular endothelium contributes to rIPC, and that inhibition of Pyk2 restores cardioprotection in T2DM through modulation of eNOS, thus limiting infarct size.

**Methods:** New Zealand Obese (NZO) mice were used as a polygenic model of T2DM. Effects of Pyk2-inhibition on endothelial function, remote ischemic preconditioning (rIPC), and infarct size (IS) after ischemia/reperfusion (I/R) were compared in NZO, eNOS KO, and C57Bl/6 (Bl6) mice. Plasma derived from mice and individuals with or without T2DM at baseline and after rIPC was transferred to isolated hearts and aortic rings to assess the effects of Pyk2-inhibition on remote tissue protection.

**Results:** Transfer experiments with plasma drawn from non-diabetic humans and mice exposed to rIPC demonstrate that endothelium-dependent signals for remote tissue protection are conveyed by plasma. Key features reflecting the glucometabolic spectrum in T2DM were detected in NZO mice, including hyperinsulinemia, insulin resistance, obesity, and impaired glucose tolerance. Similar to T2DM patients, these mice also revealed endothelial dysfunction with decreased flow-mediated dilation (FMD), reduced circulating nitrite levels, elevated arterial blood pressure, and larger infarct size after I/R. Pyk2 increased the phosphorylation of eNOS on its inhibitory site (Tyr656). Cardioprotective effects by rIPC were lost in NZO mice. Inhibition of Pyk2 restored endothelial function and rescued endothelium-dependent cardioprotection after rIPC displayed by lower IS and improved LV function post I/R.

**Conclusion:** Endothelial function contributing to remote tissue protection is severely impaired in diabetes mellitus. Proline-rich tyrosine kinase 2 is a novel target to rescue cardioprotection through endothelium-dependent remote ischemic preconditioning, advocating its role in limiting infarct size in diabetes mellitus.

**Clinical perspective What is new?:**

- Vascular endothelium contributes to remote tissue protection in ischemic preconditioning, which is severely impaired in diabetes
- Proline-rich tyrosine kinase 2 reduces eNOS-activity, causes endothelial dysfunction, and impairs cardioprotection through ischemic preconditioning
- Inhibition of proline-rich tyrosine kinase 2 restores eNOS activity, endothelial function, and cardioprotective effects of remote ischemic preconditioning limiting infarct size in an experimental model of diabetes.

**What are the clinical implications?:**

- Proper endothelial function is cirtical to maintain cardiovascular health. Endothelial dysfunction contributes to impaired remote tissue protection in diabetes.
- These data demonstrate for the first time that endothelium-dependent cardioprotection in myocardial ischemia/reperfusion through remote ischemic preconditioning can be restored in diabetes.
- Proline-rich tyrosine kinase 2 is a novel target to restore endothelium-dependent remote cardioprotection to improve the outcome of diabetic patients with acute myocardial infarction.

## Introduction

Diabetes mellitus (DM) increases the risk for myocardial infarction more than twofold ^1^. Recently, distinct phenotypes of DM that carry different cardiovascular risks have been unraveled ^2, 3^. Of these subtypes, patients with severe insulin resistance had a worse cardiovascular outcome. Patients with type 2 DM (T2DM) and acute myocardial infarction (AMI) have larger infarct sizes (IS) and impaired left ventricular (LV) function associated with distinct patterns in the myocardial texture ^4^. Therefore, especially these patients would benefit substantially from specific strategies to reduce IS after AMI ^5, 6^. One anticipated strategy represents cardioprotective effects induced by remote ischemic preconditioning (rIPC) ^7^. Neuronal, humoral, and endothelial pathways have been implicated in mediating this remote organ protection ^8^. However, translation of these experimental findings into clinical practice has been hampered so far ^9^. Remote tissue protection is lost in diabetic conditions, and strategies to reconstitute rIPC are desired while still lacking ^7, 10^.

The vascular endothelium has been implicated to contribute to rIPC. It has been suggested that endothelial nitric oxide synthase (eNOS) signaling is involved in remote tissue protection ^11^. Posttranslational mechanisms to regulate eNOS activity are numerous. One of these are different phosphorylation sites. The phosphorylation of serine (Ser) 1177 residue is well characterized to activate eNOS, whereas threonine 495 exerts an inhibitory effect on eNOS activity ^12^. Tyrosine (Tyr) phosphorylation of eNOS can act as an activator and inhibitor simultaneously: while eNOS phosphorylation on Tyr83 increases the activity ^13^, eNOS phosphorylation on Tyr657 (human sequence; Y656 mouse sequence), in contrast, strongly inhibits eNOS ^14, 15^. Tonic insulin receptor activation induces endothelial dysfunction ^14, 15^. In principle, the insulin receptor downstream signaling can induce the phosphorylation of eNOS on the stimulatory residue (Ser1177) by AKT and on the inhibitory residue (Tyr657) by Proline-rich tyrosine kinase 2 (Pyk2) ^14, 16^.

The impact of these pathways for rIPC-mediated remote organ protection in T2DM is not yet established. In this study, we provide clear evidence that endothelium-dependent cardioprotection through rIPC is lost in DM and inhibition of Pyk2 restores endothelial function, functional eNOS activity, and thus tissue protection through rIPC in DM, limiting infarct size and LV dysfunction after ischemia/reperfusion (I/R).

## Materials and Methods

A detailed description of the methods is available in the supplemental information. The data supporting the findings of this study are available from the corresponding authors upon reasonable request. Endothelial function and remote cardioprotection to influence IS after I/R was evaluated in NZO and Bl6 mice, in addition to cardiovascular characterization. As a polygenic mouse model, NZO mice are an established and well-known model resembling characteristic features of T2DM disease in humans: obesity, insulin resistance, severe hyperglycemia and disturbed glucose tolerance ^17-20^. As the prevalence of these conditions is about 60 %, each mouse was tested for manifest T2DM before the further examination.

To investigate the role of vascular endothelium in conveying signals for remote tissue protection after rIPC, we performed transfer experimets with (non)-conditioned plasma from diabetic and non-diabetic humans and mice. These were followed by in vivo and ex-vivo experiments in control and NZO mice, an experimental model of T2DM, to proof endothelial dysfunction and loss of eNOS functional activity thus impairing rIPC mediated remote tissure protection. Inhibition of Pyk2 was used to restore eNOS functional capacity and endothelium-dependent component of rIPC mediated remote tissue protection. Also in-vivo and ex-vivo control experiments with adding exogenous nitrite or scavenging of endogenous NO bioactivity were conducted.

### Patient recruitment and transfer experiments

Six patients with a medical history of cardiovascular diseases with T2DM and six age-maged controls were included. Human plasma samples were collected before and after the rIPC maneuver was performed. All clinical and hybrid experiments were performed according to the study protocol approved by the ethics committee of the Heinrich-Heine University of Duesseldorf (ID2017034183 – 5903R) and institutional guidelines. Written informed consent was obtained from all participants.

The plasma dialysates were used as a perfusion solution on murine hearts in a Langendorff-apparatus. Briefly, inotropic changes were measured by LV-end-diastolic pressure (LVEDP), contractility by the derivates of the LVEDP (+dP/dt and -dP/dt), and cardioprotective effects by the evaluation of IS (by TTC staining) following a 30 min ischemia with consecutive one-hour of reperfusion (SI Fig. 1).

Similarly, aortic rings were incubated with native and preconditioned plasma. The rings were consequently rinsed, and their function was tested in response to Ach, Phe, and SNP within a standardized protocol previously described by us ^21^.

### Animals

Animal experiments were approved by the local animal ethics committee (LANUV, NRW, Germany) and performed according to the European Convention for the Protection of Vertebrate Animals used for Experimental and other Scientific Purposes (Council of Europe Treaty Series No. 123). Animal care followed institutional guidelines, and experimental planning and execution followed the ARRIVE recommendations ^22^. All experiments were performed in 15-18 week-old New Zealand Obese (NZO) mice (NZO/HILtJ) ^23^, C57BL/6J (Bl6), or global eNOS knockout (eNOS^-/-^) ^24^ mice.

### Pyk2 inhibition

Pharmacological Pyk2 inhibition was achieved with PF-431396 hydrate (Sigma-Aldrich, EUA), which was administrated (5 μg/g, i.p.) at least 15 minutes before the respective measurement or performed protocol, as previously described ^25^.

### Ultrasound imaging measurements

High-resolution micro-ultrasound imaging measurements (echocardiography, FMD, PWV) were performed on Vevo 2100/3100 (Visual Sonics Inc., Toronto, Canada). Mice were kept in a stable cardiopulmonary state by isoflurane anesthesia (1.5-2%). Analysis of the cardiac function (18-38 MHz transducer) was performed as described ^21^. It included measurements of left ventricular (LV) end-systolic (ESV) and end-diastolic (EDV) volumes, left ventricular ejection fraction (EF), cardiac output (CO), and stroke volume (SV), which was normalized to body surface area and body weight.

Vascular- and endothelial function measurements (30-70 MHz transducer) relied on a baseline recording of the diameter of the A. iliaca externa followed by five-minute occlusion of the artery distally of the transducer. The maximal dilation of the vessel in response to reperfusion was evaluated as FMD ^21^. Vascular stiffness was assessed as pulse wave velocity (PWV) in the common carotid artery by dividing the distance (ΔD) of the blood flow by the time (ΔT) as described ^21^.

### Remote ischemic preconditioning

The rIPC protocol consisted of four cycles of ischemic preconditioning (IPC, Figure SI1). Anesthetized mice (1.5-2% isoflurane) were placed with a vascular occluder around their left hindlimb, which was inflated to a pressure of 200 mmHg and kept constant for five minutes. Subsequently, the occluder was deflated, allowing tissue reperfusion for five minutes. This protocol was repeated four times before further measurements or procedures were implemented. The same protocol was performed in humans; a blood pressure cuff was placed at a forearm following this protocol.

### Model of ischemia-reperfusion and assessment of infarct sizes

After adequate anesthesia with buprenorphine (0.1 mg/kg), mice were intubated and kept on isoflurane anesthesia (induction 3% V/V and maintenance 2% V/V) via a rodent ventilator side port. Regardless of the protocol used, all groups were exposed to the exact duration of anesthesia. The thorax was opened, and local myocardial ischemia was induced by the occlusion of the left descendent artery (LAD) for 30 minutes. ST elevations confirmed the successful occlusion. Consequently, the LAD was reperfused, the thorax was closed, and rodents were subjected to 24 hours of reperfusion, following a standardized protocol ^26^. On the following day, infarct sizes (IS) were determined by triphenyl tetrazolium chloride (TTC) staining, as described ^26^. IS was expressed as a percentage of the infarcted tissue area within the total area at risk (AAR).

### Statistical analysis

The sample size was calculated prior by using G-Power V3.1. (Heinrich Heine University of Duesseldorf). Results are displayed as mean ± standard error of the mean (SEM). For repeated measurements, data were analyzed by one-way and two-way analysis of variance (ANOVA), as appropriate, followed by Bonferroni’s or Sidak’s post hoc tests. Where indicated, an unpaired Student’s t-test was used to determine whether the two data groups differed significantly. Normal distribution was tested through the D’Agostino-Pearson test. *p* < 0.05 was considered statistically significant. Statistical analysis was done using GraphPad Prism 9 for Windows (GraphPad Software, San Diego, CA, USA).

## Results

### In patients with T2DM, plasma lacks signals conveying remote tissue protection

To test the hypothesis that rIPC induces the release of circulating messengers to promote remote tissue protection conveyed in the circulation, samples of plasma drawn after repetitive limb ischemia were transferred to Langendorff hearts subjected to ischemia/reperfusion or isolated aortic rings (Figure 1A). When perfused with the preconditioned plasma samples from healthy participants, the isolated hearts showed a better LV hemodynamics and diminished IS after global ischemia (Figure 1B). Aortic rings showed an improved endothelium-dependent dilation after exposure to healthy preconditioned plasma (Figure 1C). These transfer experiments demonstrate that in patients without T2DM, rIPC induced the release of circulating messengers into the plasma, which conveyed cardioprotection. These effects on both hearts and aortic rings were entirely lost in preconditioned plasma derived from patients with T2DM, demonstrating that the signal origin, the endothelium, was dysfunctional. Smooth muscle dilatory response was not altered in aortic ring segments preincubated with conditioned plasma from either diebetic or non-diabetic humans (Figure SI 6A). Our results provide evidence that rIPC-mediated remote protective signaling can be conveyed through plasma components, which are abrogated under diabetic conditions. The subjects’ full demographic data are summarized in SI Table 1.

**Figure 1:**
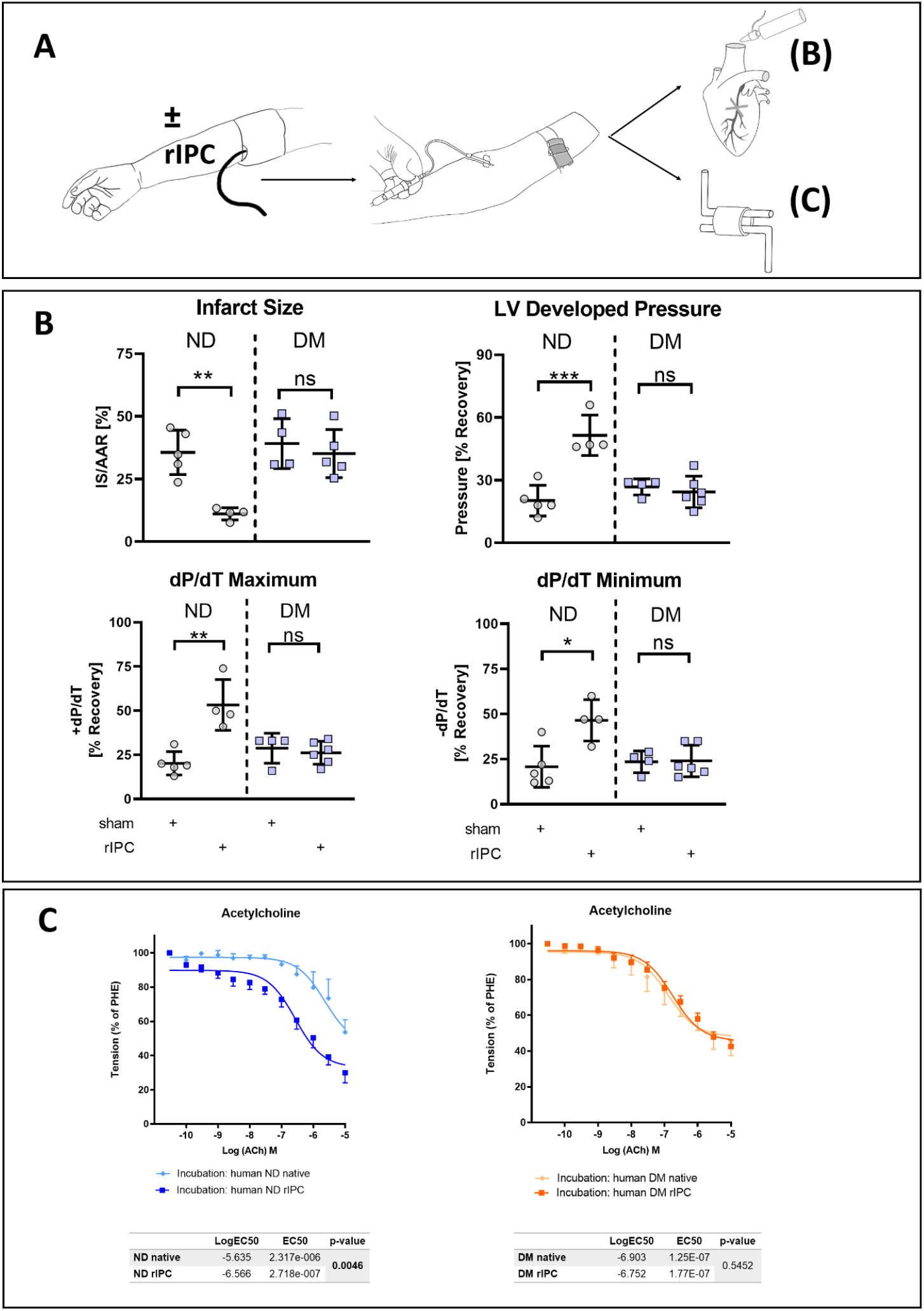
Remote tissue protection is abolished in patients with T2DM. **(A)** Remote ischemic conditioning was performed at the forearm of individuals with (T2DM – diabetic patients) or without T2DM (ND – non-diabetic patients). After blood collection, plasma was transfered to either Langendorff hearts prior to global ischemia or aortic rings from control mice. **(B)** rIPC-treated plasma from patients without T2DM but not the plasma from patients with T2DM decreased infarct size (IS) per area at risk (AAR), optimized left ventricular developed pressure, cardiac relaxation (dP/dTmin) and contractility (dP/dTmax). **(C)** Endothelium-dependent dilatory response in isolated rings was significantly improved with rIPC-conditioned plasma from individuales without T2DM but not from patients with T2DM. n = 5-6, * p < 0.05; ** p < 0.01; *** p < 0.001.

### Cardiovascular and metabolic characteristics of NZO mice

Circulating insulin levels were elevated, glucose levels were higher during the glucose tolerance test, and body weight was increased in NZO mice, proving all hallmarks of the glucometabolic spectrum of T2DM (Figure 2A). While body-weight was increased, the heart-weight to body-weight ratios were similar compared to Bl6 mice (Figure 2A). Similar results were found in cardiac functional parameters, as evaluated by echocardiography (SI Table 2). LV functional parameters in terms of stroke volume (SV) and cardiac output (CO) ejection fraction (EF) were similar in NZO mice as compared to Bl6 mice. Pyk2-inhibition did not affect these parameters in either of the strains at baseline (SI Table 2). The impact of T2DM on systemic hemodynamics was evaluated with millar catheterization. NZO mice displayed a significantly higher systolic- and mean arterial pressure with a consecutive rise in total peripheral resistance (Figure 2B). The application of Pyk2 inhibitor reversed the increase in arterial blood pressure in NZO mice (Figure 2B, SI Table 2).

**Fig 2:**
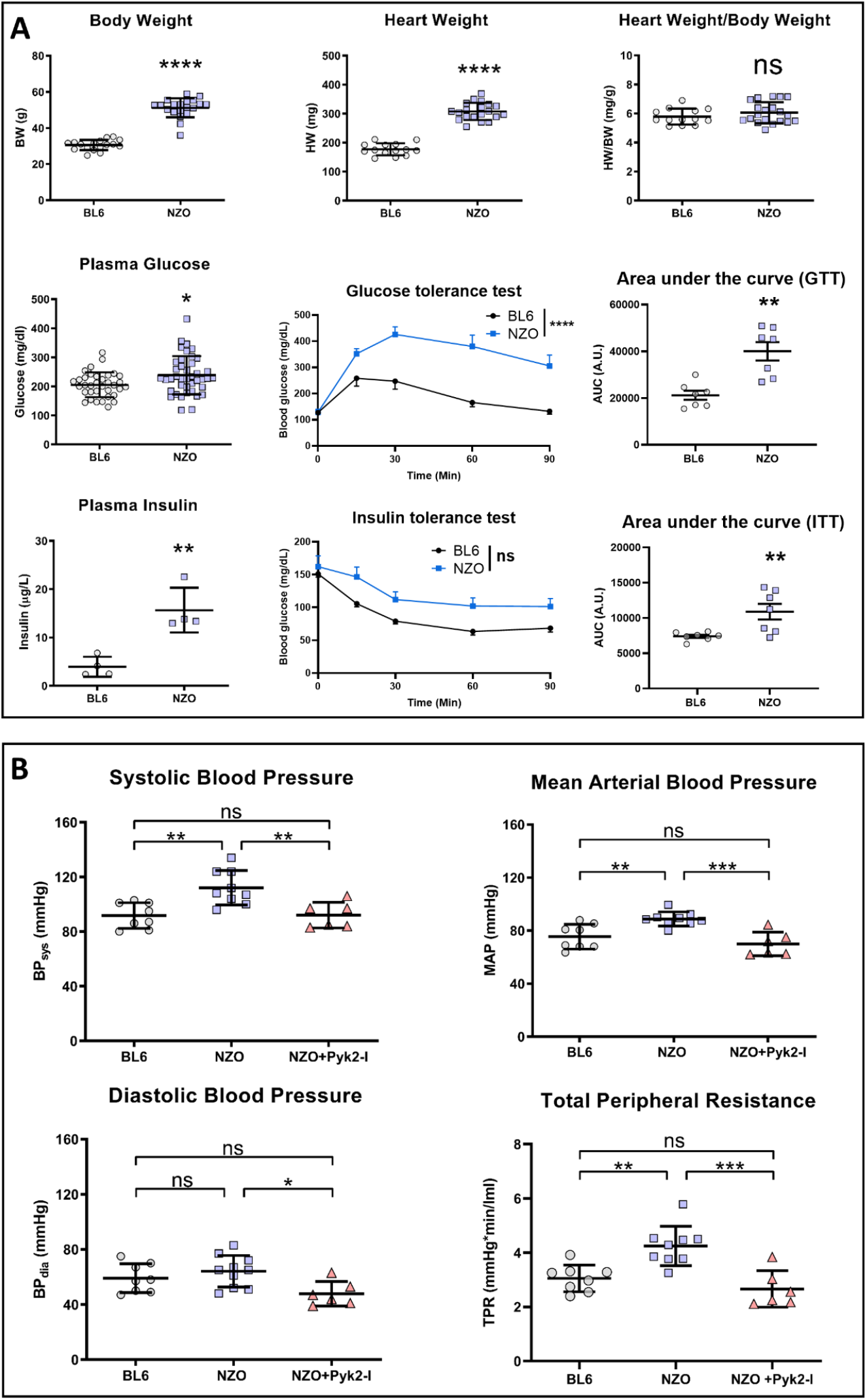
Vascular and metablic alterations in NZO-mice. **(A)** Body weight; heart weight, and heart weight/body weight (HW/BW) ratio of Bl6 and NZO mice. Blood glucose levels Bl6 and NZO mice; plasma insulin levels; area under the curve (AUC) of the blood glucose during the glucose tolerance test (GTT); glucose tolerance test (GTT); plasma insulin levels during the glucose tolerance test. **(B)** Systolic pressure, mean arterial blodd pressure (MAP), and total peripheral resistance (TPR) was significantly increased in NZO as compared to control mice. The inhibition of Pyk2 normalized blood pressure and vascular resistance in NZO mice as compared to BL6 mice. n = 14-20 for A) and 6 for B-C); * p < 0.05; ** p < 0.01; **** p < 0.0001.

In-vivo (FMD) and ex-vivo (organ bath) experiments were performed to assess endothelium-dependent vasodilation. FMD response was abolished in NZO mice and preserved in Bl6 mice (Figure 3A). Due to the lack of endothelial response in the FMD of eNOS knockout (KO) mice and in Bl6 mice treated with pharmacological eNOS inhibitor LNAME, we validate eNOS dependent components of FMD after limb ischmia (SI Figure 2). Similar results were found ex-vivo in organ bath experiments (Figure 3B and SI Figure 3B). Endothelial dysfunction has been shown to amplify arterial stiffness ^27^. The elasticity of conductive arteries in NZO mice was severely impaired, as indicated by increased pulse wave velocity (PWV) (Figure 3C). This was comparable to PWV observed in eNOS^-/-^ mice (not shown), denoting the importance of a functional eNOS for preserved arterial vessel elasticity. To summarize, NZO mice exhibited metabolic key features of T2DM, increased arterial stiffness and blood pressure, severe endothelial dysfunction, and preserved cardiac function at baseline (SI Table 2).

**Figure 3:**
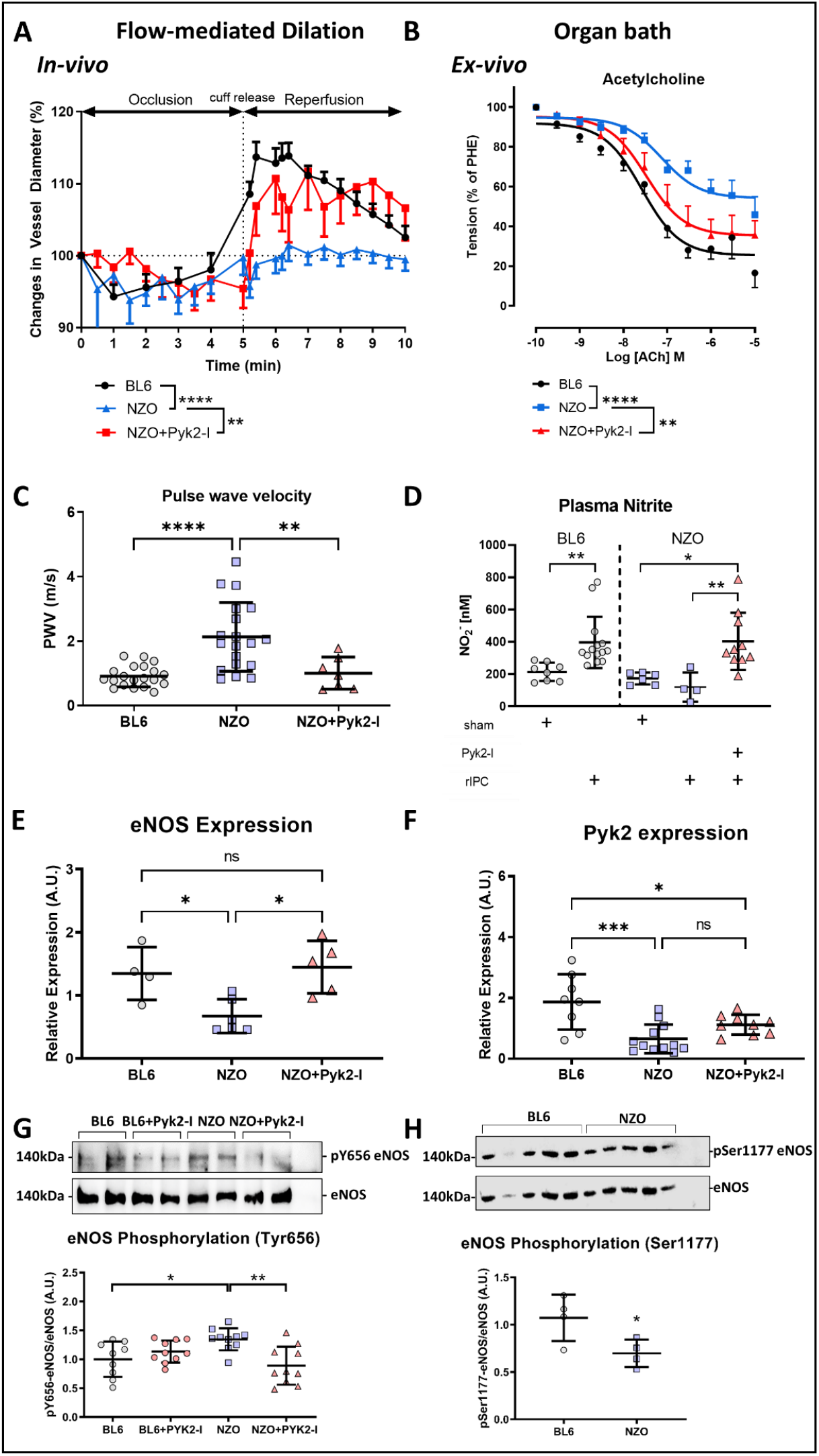
Pyk2-inhibition restores eNOS activity in T2DM. **(A)**Endothelial function assessed in-vivo by non-invasive measurement of flow-mediated dilation (FMD). Pyk2-inhibition rescues manifest endothelial dysfunction in diabetic NZO mice. **(A)**Organ-bath experiments revealed a significant improvement of endothelial function in response to Ach after Pyk2-inhibition in NZO mice. **(C)** Pyk2-inhibition normalizes pulse-wave-velocity (PWV) in NZO mice. **(D)** Increased plasma nitrite levels after Pyk2-inhibition reveal a reactivation of eNOS functional capacity after shear-mediated activation by rIPC. **(E, F)** Gene expression of eNOS and Pyk2 in NZO and BL6 mice hearts. **G, H)** Representative western blots and densitometric analysis of eNOS, key activation site (Ser1177), and inhibition-status (Tyr656) in Bl6 and NZO mice. **(A – D):** n = 10-15; * p < 0.05; ** p < 0.01; *** p < 0.001; **** p < 0.0001. **(D – H):** n = 4-8; * p < 0.05; ** p < 0.01; **** p < 0.0001.

### Inhibition of Pyk2 regains eNOS activity and endothelial function

Changes in FMD imply alterations in the activity status of eNOS, expressed by its phosphorylation and in plasma nitrite levels (as a product of NO synthesis). To test the impact of Pyk2-inhibition on the functional status (FMD, organ bath) and activation levels (phosphorylation, plasma nitrite levels) of eNOS, we performed several inv-vivo and ex-vivo experiments in Bl6 and NZO mice, both with and without Pyk2-inhibiton..

Pyk2-inhibition significantly improved endothelium-dependent relaxation of NZO mice, both in-vivo measured by the FMD and ex-vivo by aortic ring assay (Figure 3B), strengthening the notion that Pyk2-inhibition restores eNOS functionality. In contrast, Pyk2-inhibition did not change endothelial function in Bl6 mice (SI Figure 3A). The inhibitor did not affect the VSMC layer (in response to Phe and SNP) in any strains (SI Figure 3B, C). Vehicle control (DMSO) did neither affect the dose-response curves nor in-vivo FMD results (data not shown).

NZO mice showed a higher phosphorylation of the inhibitory site of eNOS on Tyr656, accompanied by lower phosphorylation levels of the primary activation site Ser1177 (Figure 3G and 3H). Moreover, gene expression of Pyk2 and eNOS was markedly reduced in NZO mice (Figure 3E and 3F). Inhibition of Pyk2 diminished the phosphorylation levels of eNOS on its inhibitory site (Tyr656) in NZO mice but did not change phosphorylation levels in Bl6 mice (Figure 3G). Expression of eNOS was decreased at baseline and normalized after Pyk2-inhibition in NZO mice (Figure 3D), implying a counter regulatory mechanism of Pyk2. Plasma levels of nitrite (NO ^-^) increased after applying the rIPC maneuver in Bl6 but not in NZO mice (Figure 3E). Inhibition of Pyk2 restores eNOS functional capacity in NZO mice to be activated by shear stress, as demonstrated by increased levels of nitrite (Figure 3D). These results demonstrate the potential of Pyk2-inhibition to restore eNOS functional activity.

### Endothelium-dependent remote cardioprotection is impaired in NZO mice and reconstituted by Pyk2-inhibition

In NZO mice, Pyk2-inhibition reconstitute endothelial function and restores eNOS functional activity. Next, we aimed to analyze endothelium-dependent components of rIPC mediated cardioprotection in non-diabetic NZO and Bl6 mice and to investigate its impact on IS and LV functional parameters after I/R.

Following I/R, NZO mice suffered from larger infarct sizes and more significant functional impairment of LV function than Bl6 mice (Figure 4 A and B). RIPC significantly reduced IS and improved LV function in Bl6 but not in NZO-hearts after I/R. Supplementation of nitrite did not lead to an additional reduction of IS in Bl6 mice (compared to hearts receiving rIPC) but significantly reduced IS and improved LV function in NZO mice. Scavenging of circulatin NO bioactivity through CPTIO severely inflicted I/R damage to BL6 hearts, but it did not affect IS or LV functional parameters in NZO mice (Figure 4 A and B, Table 1). RIPC maneuver with simultaneous administration of CPTIO neutralizes cardioprotective effects, as illustrated by the increase of IS and functional parameters. To summarize, these results support the notion that remote cardioprotection by rIPC is, at least in part, mediated by circulating NO bioactivity derived from endothelial eNOS (Figure 4 A and B, Table 1) and that these beneficial effects are severly disturbed in T2DM.

**Figure 4:**
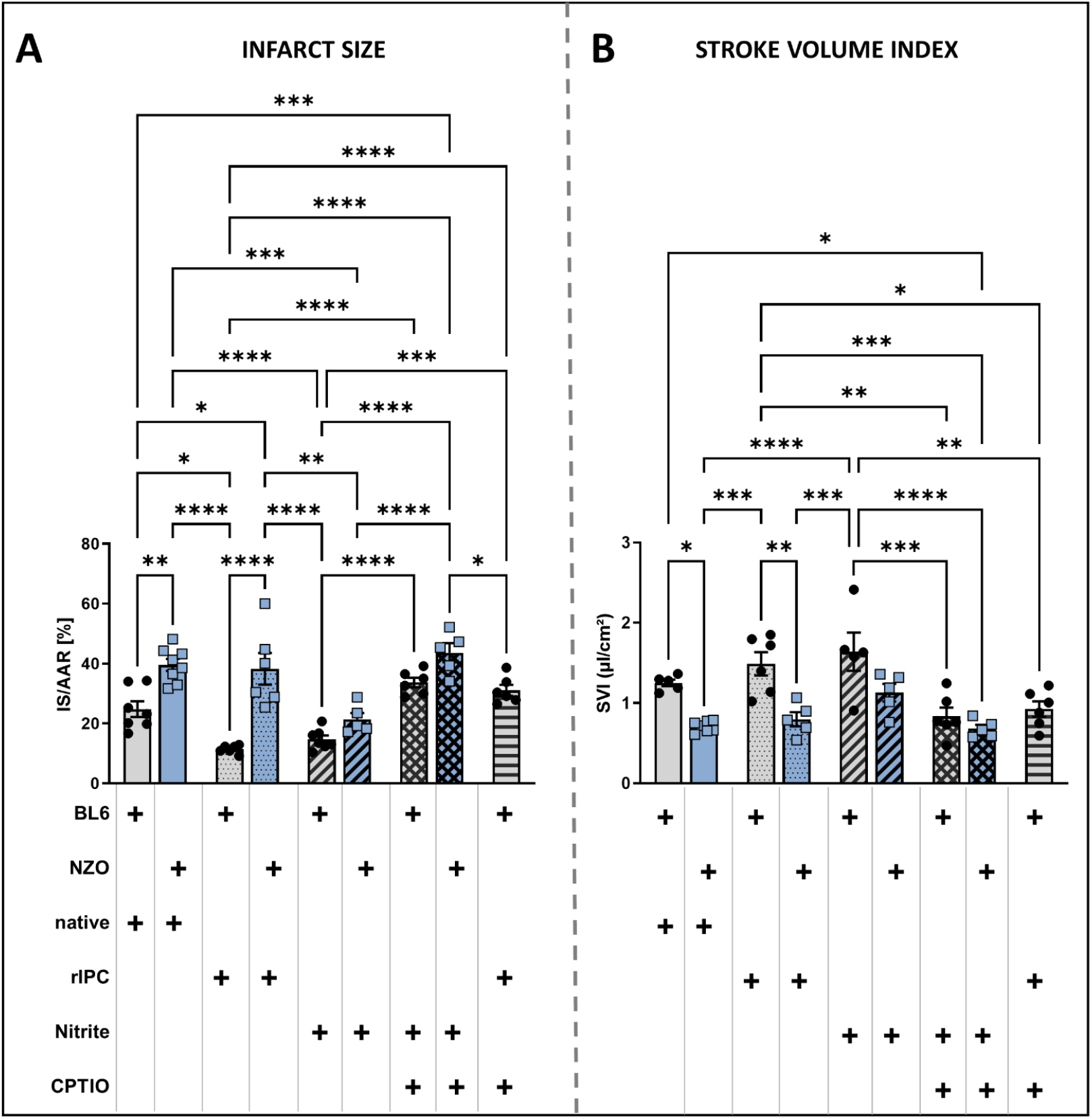
Impaired endothelium-dependent remote cardioprotection in T2DM. Evaluation of infarct sizes (IS) per area at risk (AAR) in distinct protocols (please refer to SI Figure 3 for detailed information on protocols). Evaluation of LV functional parameters was performed 24h after I/R by echocardiography. Specific intervention per group (control mice in greay bars, and NZO mice in blue bars) is highlighted with a plus sign. Sham mice received no treatment before ligation of the left descending artery. Groups treated with the rIPC maneuver were admitted to repetitive phases of hindlimb ischemia immediately before LAD ligation. Nitrite or cPTIO administration was performed 5 minutes before LAD ligation. **(A)** Maneuver of rIPC significantly reduces IS in Bl6-mice and improved LV Function **(B)**, while this cadioportective effect was los in NZO mice. Nitrite, as an exobegnous reservoir of NO bioactivity was neutral in control mice, but significantly reduced IS in NZO mice. Co-administration of the NO scavenger cPTIO selectively neutralizes the cardioprotective effects of rIPC in control mice and nitrite in NZO mice. **(B)** Stroke-Volume index corresponding to the distinct intervention group and changes in IS as depicted in **(A)**. n = 6-8; * p < 0.05; ** p < 0.01; *** p < 0.001; **** p < 0.0001.

**Table 1:**
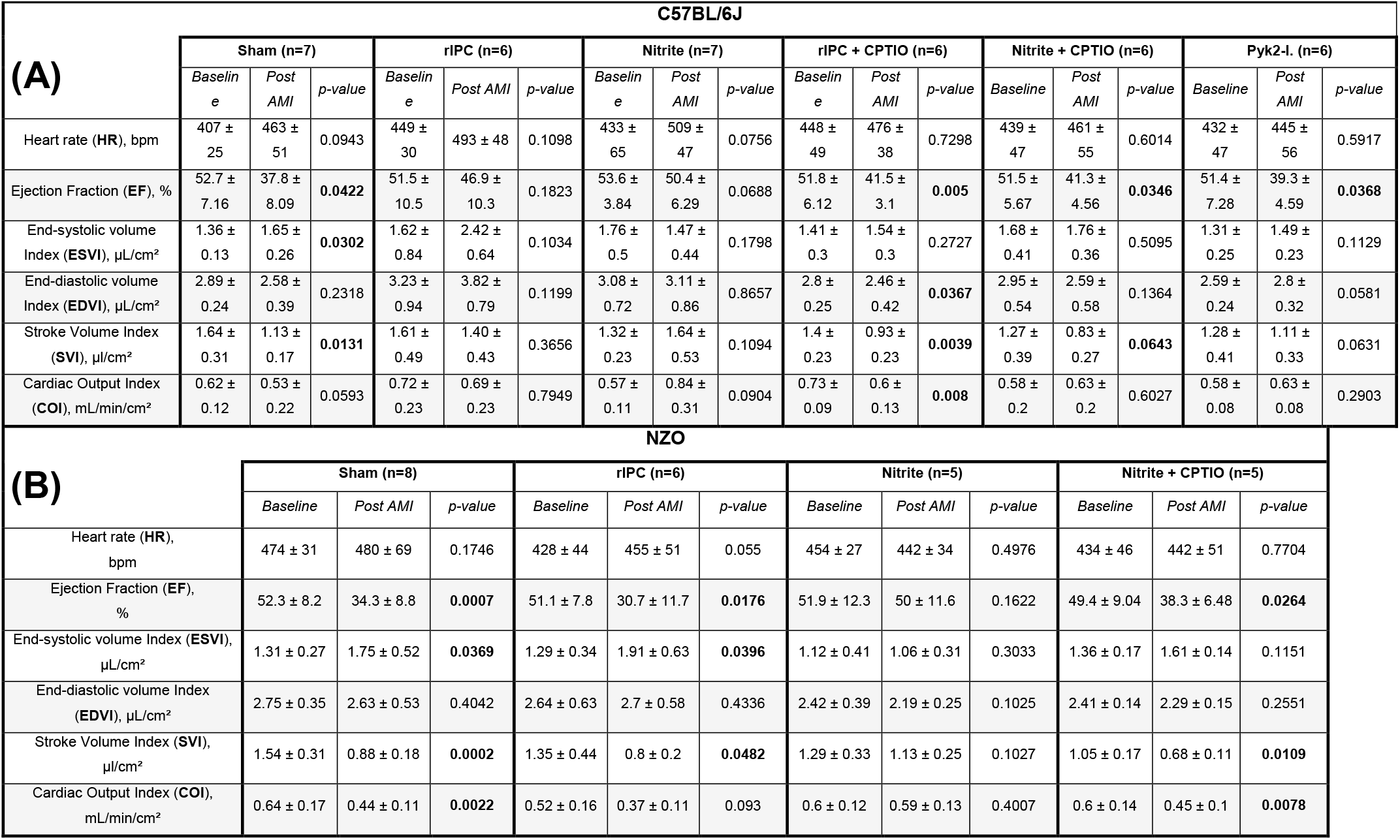
Echocardiographic parameters of mice subjected to AMI and corresponding IS are shown in Figure 4 (A) and (B).

| (A) | C57BL/6J |  |  |  |  |  |  |  |  |  |  |  |  |  |  |  |  |  |
| --- | --- | --- | --- | --- | --- | --- | --- | --- | --- | --- | --- | --- | --- | --- | --- | --- | --- | --- |
|  | Sham (n=7) |  |  | rIPC (n=6) |  |  | Nitrite (n=7) |  |  | rIPC + CPTIO (n=6) |  |  | Nitrite + CPTIO (n=6) |  |  | Pyk2-I. (n=6) |  |  |
|  | Baseline | Post AMI | p-value | Baseline | Post AMI | p-value | Baseline | Post AMI | p-value | Baseline | Post AMI | p-value | Baseline | Post AMI | p-value | Baseline | Post AMI | p-value |
| Heart rate (HR), bpm | 407 ± 25 | 463 ± 51 | 0.0943 | 449 ± 30 | 493 ± 48 | 0.1098 | 433 ± 65 | 509 ± 47 | 0.0756 | 448 ± 49 | 476 ± 38 | 0.7298 | 439 ± 47 | 461 ± 55 | 0.6014 | 432 ± 47 | 445 ± 56 | 0.5917 |
| Ejection Fraction (EF), % | 52.7 ± 7.16 | 37.8 ± 8.09 | **0.0422** | 51.5 ± 10.5 | 46.9 ± 10.3 | 0.1823 | 53.6 ± 3.84 | 50.4 ± 6.29 | 0.0688 | 51.8 ± 6.12 | 41.5 ± 3.1 | **0.005** | 51.5 ± 5.67 | 41.3 ± 4.56 | **0.0346** | 51.4 ± 7.28 | 39.3 ± 4.59 | **0.0368** |
| End-systolic volume Index (ESVI), $\mu\text{L}/\text{cm}^2$ | 1.36 ± 0.13 | 1.65 ± 0.26 | **0.0302** | 1.62 ± 0.84 | 2.42 ± 0.64 | 0.1034 | 1.76 ± 0.5 | 1.47 ± 0.44 | 0.1798 | 1.41 ± 0.3 | 1.54 ± 0.3 | 0.2727 | 1.68 ± 0.41 | 1.76 ± 0.36 | 0.5095 | 1.31 ± 0.25 | 1.49 ± 0.23 | 0.1129 |
| End-diastolic volume Index (EDVI), $\mu\text{L}/\text{cm}^2$ | 2.89 ± 0.24 | 2.58 ± 0.39 | 0.2318 | 3.23 ± 0.94 | 3.82 ± 0.79 | 0.1199 | 3.08 ± 0.72 | 3.11 ± 0.86 | 0.8657 | 2.8 ± 0.25 | 2.46 ± 0.42 | **0.0367** | 2.95 ± 0.54 | 2.59 ± 0.58 | 0.1364 | 2.59 ± 0.24 | 2.8 ± 0.32 | 0.0581 |
| Stroke Volume Index (SVI), $\mu\text{L}/\text{cm}^2$ | 1.64 ± 0.31 | 1.13 ± 0.17 | **0.0131** | 1.61 ± 0.49 | 1.40 ± 0.43 | 0.3656 | 1.32 ± 0.23 | 1.64 ± 0.53 | 0.1094 | 1.4 ± 0.23 | 0.93 ± 0.23 | **0.0039** | 1.27 ± 0.39 | 0.83 ± 0.27 | **0.0643** | 1.28 ± 0.41 | 1.11 ± 0.33 | 0.0631 |
| Cardiac Output Index (COI), $\text{mL}/\text{min}/\text{cm}^2$ | 0.62 ± 0.12 | 0.53 ± 0.22 | 0.0593 | 0.72 ± 0.23 | 0.69 ± 0.23 | 0.7949 | 0.57 ± 0.11 | 0.84 ± 0.31 | 0.0904 | 0.73 ± 0.09 | 0.6 ± 0.13 | **0.008** | 0.58 ± 0.2 | 0.63 ± 0.2 | 0.6027 | 0.58 ± 0.08 | 0.63 ± 0.08 | 0.2903 |
| (B) | NZO |  |  |  |  |  |  |  |  |  |  |  |  |  |  |  |  |  |
|  | Sham (n=8) |  |  | rIPC (n=6) |  |  | Nitrite (n=5) |  |  | Nitrite + CPTIO (n=5) |  |  |  |  |  |  |  |  |
|  | Baseline | Post AMI | p-value | Baseline | Post AMI | p-value | Baseline | Post AMI | p-value | Baseline | Post AMI | p-value | Baseline | Post AMI | p-value | Baseline | Post AMI | p-value |
| Heart rate (HR), bpm | 474 ± 31 | 480 ± 69 | 0.1746 | 428 ± 44 | 455 ± 51 | 0.055 | 454 ± 27 | 442 ± 34 | 0.4976 | 434 ± 46 | 442 ± 51 | 0.7704 |  |  |  |  |  |  |
| Ejection Fraction (EF), % | 52.3 ± 8.2 | 34.3 ± 8.8 | **0.0007** | 51.1 ± 7.8 | 30.7 ± 11.7 | **0.0176** | 51.9 ± 12.3 | 50 ± 11.6 | 0.1622 | 49.4 ± 9.04 | 38.3 ± 6.48 | **0.0264** |  |  |  |  |  |  |
| End-systolic volume Index (ESVI), $\mu\text{L}/\text{cm}^2$ | 1.31 ± 0.27 | 1.75 ± 0.52 | **0.0369** | 1.29 ± 0.34 | 1.91 ± 0.63 | **0.0396** | 1.12 ± 0.41 | 1.06 ± 0.31 | 0.3033 | 1.36 ± 0.17 | 1.61 ± 0.14 | 0.1151 | | | | | | |
| End-diastolic volume Index (EDVI), $\mu\text{L}/\text{cm}^2$ | 2.75 ± 0.35 | 2.63 ± 0.53 | 0.4042 | 2.64 ± 0.63 | 2.7 ± 0.58 | 0.4336 | 2.42 ± 0.39 | 2.19 ± 0.25 | 0.1025 | 2.41 ± 0.14 | 2.29 ± 0.15 | 0.2551 | | | | | | |
| Stroke Volume Index (SVI), $\mu\text{L}/\text{cm}^2$ | 1.54 ± 0.31 | 0.88 ± 0.18 | **0.0002** | 1.35 ± 0.44 | 0.8 ± 0.2 | **0.0482** | 1.29 ± 0.33 | 1.13 ± 0.25 | 0.1027 | 1.05 ± 0.17 | 0.68 ± 0.11 | **0.0109** | | | | | | |
| Cardiac Output Index (COI), $\text{mL}/\text{min}/\text{cm}^2$ | 0.64 ± 0.17 | 0.44 ± 0.11 | **0.0022** | 0.52 ± 0.16 | 0.37 ± 0.11 | 0.093 | 0.6 ± 0.12 | 0.59 ± 0.13 | 0.4007 | 0.6 ± 0.14 | 0.45 ± 0.1 | **0.0078** | | | | | | |

In-vivo experiments in both strains aimed to validate our ex-vivo results. Indeed, targeting Pyk2 by its inhibition restored rIPC-dependent remote cardioprotection in NZO mice, demonstrated by their significantly reduced IS and improved LV functional parameters (Figure 5A and B, Table 2). CPTIO neutralized all protective effects, validating again eNOS-metabolites to convey signals for heterocellular cardioprotection by rIPC. As rIPC-dependent remote protection can be mediated by neuronal pathways as well ^7^, we tested the impact of the neuronal pathway in NZO and Bl6 mice by dissecting the femoral nerve prior rIPC maneuver (SI Figure 5). RIPC partially conferred protection displayed by slightly lower IS, while the extent of cardioprotection compared to rIPC alone. This demonstrated, that both endothelium-dependent signaling and neuronal pathways contribute to remote tissure protection through rIPC.

**Figure 5:**
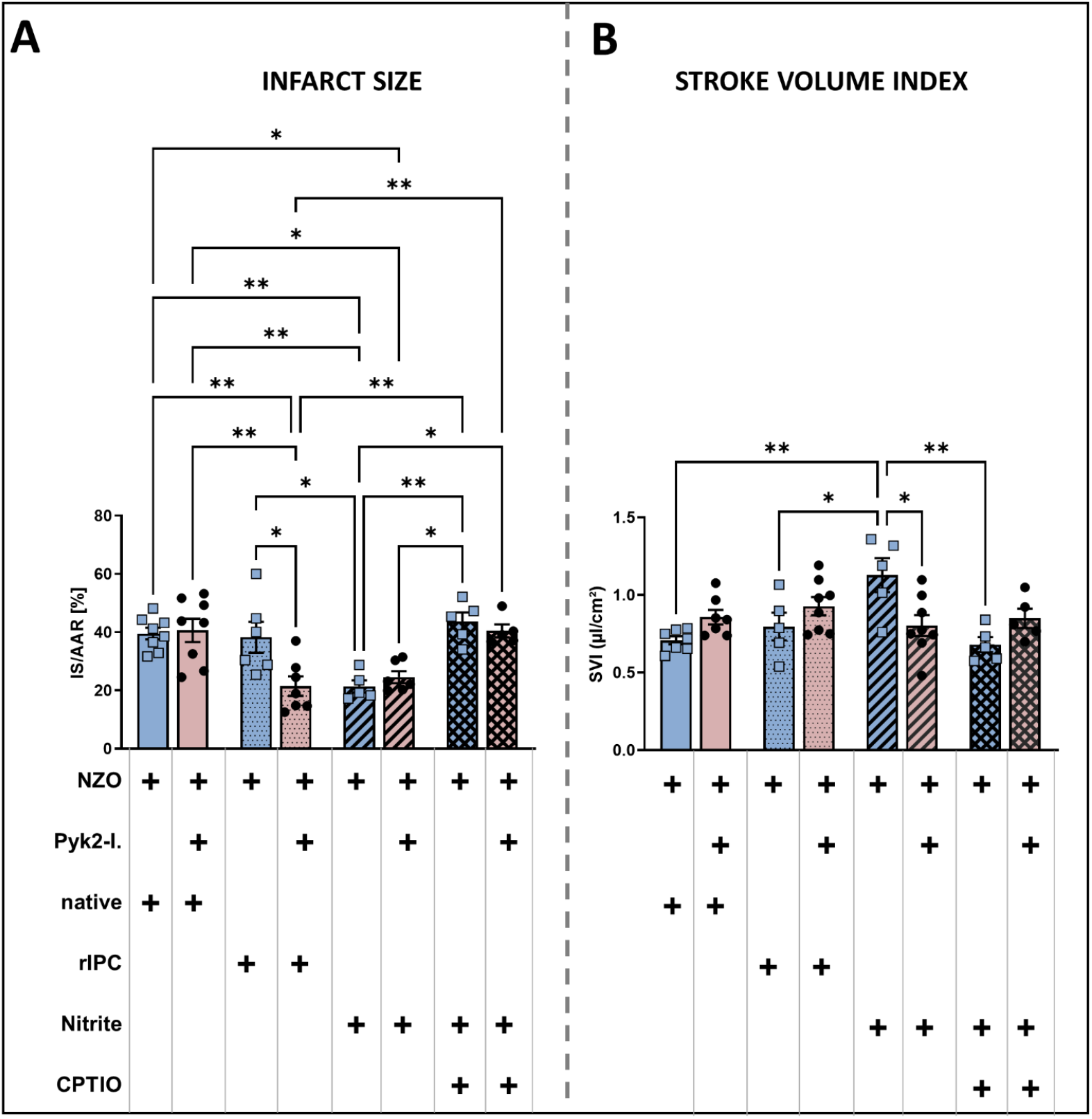
Pyk2-inhibition restores endothelium-dependent remote cardioprotection in T2DM. Evaluation of infarct sizes (IS) per area at risk (AAR) in distinct protocols (please refer to SI Figure 3 for detailed information on protocols). Evaluation of stroke volume (SV) was performed 24h after I/R by echocardiography. Distinct intervention per group is highlighted with a plus sign. Pyk2-inhibitor (Pyk2-I) was injected 15 minutes before different protocols. **(A)** In NZO mice Pyk2-inition restores cardioprotective effects induced by the rIPC maneuver, indicated by smaller IS and improved SV. Administration of NO-scavenger cPTIO neutralizes this cardioprotective effect, indicating that eNOS contributed to rIPC-dependent cardioprotection. Exogenous nitrite was cardioprotective, but neutral on top of rIPC. **(B)** Stroke-Volume index corresponding to the distinct intervention group. n = 6-8; * p < 0.05; ** p < 0.01; *** p < 0.001; **** p < 0.0001.

**Table 2:** Echo parameters of mice subjected to AMI, corresponding IS are shown in

|  | NZO + Pyk2-Inhibitor |  |  |  |  |  |  |  |  |  |  |  |
| --- | --- | --- | --- | --- | --- | --- | --- | --- | --- | --- | --- | --- |
|  | Sham (n=8) |  |  | rIPC (n=7) |  |  | Nitrite (n=6) |  |  | rIPC + CPTIO (n=5) |  |  |
|  | Baseline | Post AMI | p-value | Baseline | Post AMI | p-value | Baseline | Post AMI | p-value | Baseline | Post AMI | p-value |
| Heart rate (HR),<br>bpm | 459 ± 41 | 475 ± 35 | 0.4778 | 456 ± 27 | 440 ± 49 | 0.4036 | 411 ± 72 | 478 ± 28 | 0.0735 | 415 ± 54 | 449 ± 51 | 0.0577 |
| Ejection Fraction (EF),<br>% | 51.4 ± 8.2 | 42.2 ± 11.6 | **0.0244** | 53.9 ± 12.8 | 54.6 ± 11.1 | 0.9165 | 58.4 ± 11.4 | 52 ± 9.1 | **0.0152** | 51.8 ± 5.69 | 38.2 ± 3.13 | **0.0148** |
| End-systolic volume Index (ESVI),<br>μL/cm<sup>2</sup> | 1.26 ± 0.29 | 1.24 ± 0.14 | 0.5616 | 1.11 ± 0.41 | 0.88 ± 0.21 | 0.3383 | 0.73 ± 0.26 | 0.73 ± 0.19 | 0.9743 | 1.06 ± 0.44 | 1.15 ± 0.18 | 0.8737 |
| End-diastolic volume Index (EDVI),<br>μL/cm<sup>2</sup> | 2.42 ± 0.29 | 2.08 ± 0.15 | **0.0332** | 2.38 ± 0.37 | 1.98 ± 0.34 | 0.1474 | 1.79 ± 0.26 | 1.53 ± 0.26 | 0.2123 | 1.88 ± 0.4 | 2.11 ± 0.29 | 0.8026 |
| Stroke Volume Index (SVI),<br>μL/cm<sup>2</sup> | 1.57 ± 0.12 | 0.91 ± 0.20 | **0.0092** | 1.37 ± 0.31 | 1.1 ± 0.35 | 0.327 | 1.36 ± 0.32 | 1.18 ± 0.19 | 0.1296 | 1.22 ± 0.13 | 0.85 ± 0.18 | 0.0535 |
| Cardiac Output Index (COI),<br>mL/min/cm<sup>2</sup> | 0.64 ± 0.09 | 0.63 ± 0.15 | 0.9098 | 0.58 ± 0.14 | 0.48 ± 0.16 | 0.2222 | 0.43 ± 0.11 | 0.39 ± 0.08 | 0.3062 | 0.64 ± 0.84 | 0.42 ± 0.04 | 0.4223 |

To specifically validate our in-vivo experiments on the impact of eNOS functionality on circulating messengers to convey cardioprotection, we tested the reconstitution of the tissue protective effect in ex-vivo experiments transferring plasma from non-diabetic and diabetic mice after rIPC pretreated with the Pyk2-inhibitor or not (Figure 6A). Endothelium-dependent relaxation in response to acetylcholine, and carbachol as control, was selectively improved in aortic rings preincubated with conditioned plasma obtained from NZO mice pretreated with the Pyk2 inhibitor (Figure 6B). These data provide unequivocal evidence that Pyk2-inhibition reconstitutes circulating messengers to confer for remote tissue protection in diabetic conditions.

**Figure 6:**
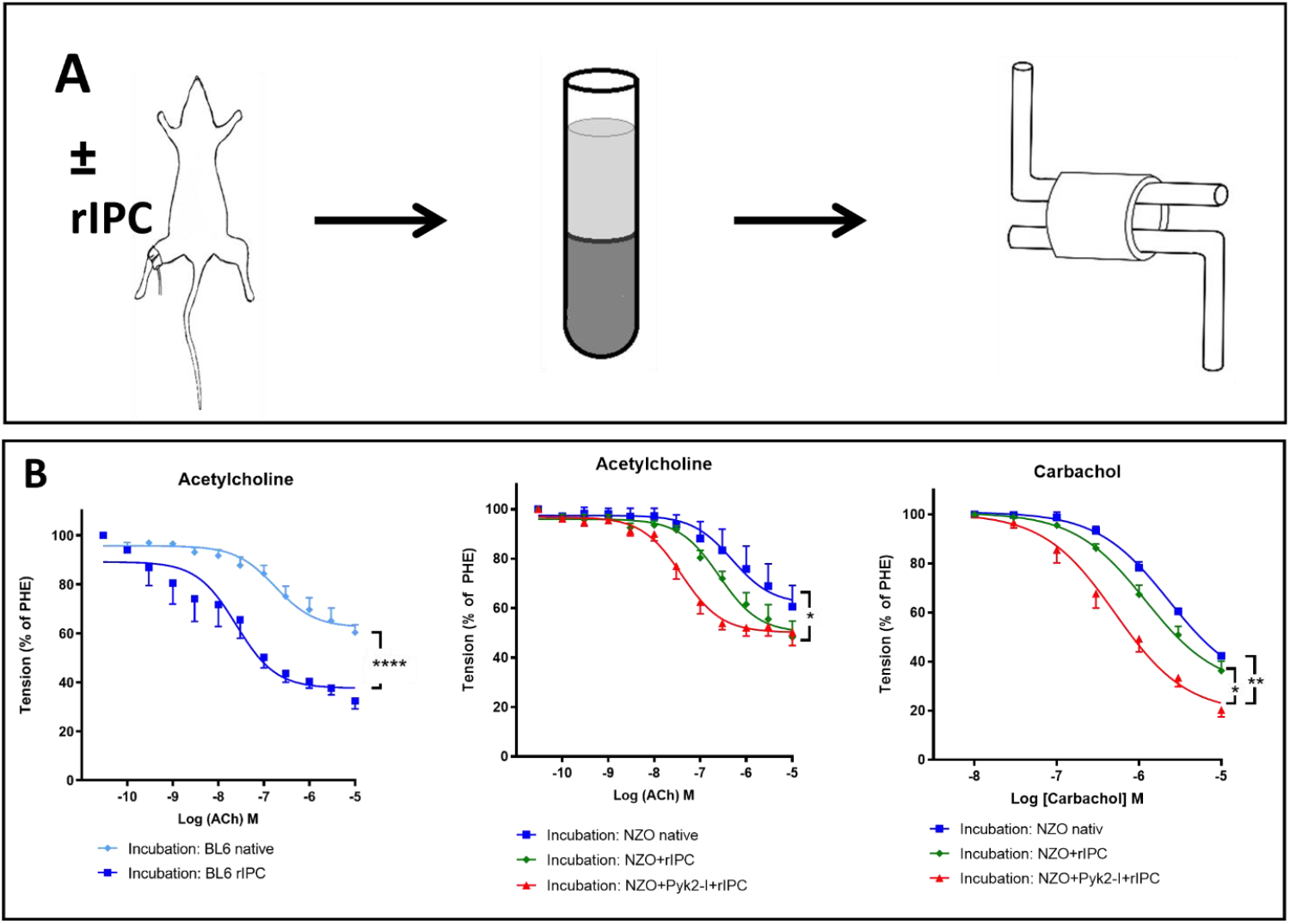
Pyk2-inhibition restores eNOS activity to induce rIPC associated release of circulating messengers into plasma for remote tissue protection. **(A)** RIPC maneuver was performed on Bl6 or NZO mice with or without Pyk2-inhibition. Blood samples were drawn, and plasma was transferred to aortic rings of Bl6 mice in organ baths. **(B)** Conditioned plasma originating from control mice improved endothelial function in aortic rings, while plasma from NZO mice treated with rIPC show now effect. Exposing NZO mice to rIPC in the presence of the Pyk2-inhibitor, their conditioned plasma improved endothelial function in aortic rings. Complementary results to organ-bath experiments as shown in Figure 1 C. n = 6-8; ** p < 0.01; **** p < 0.0001.

## Discussion

The present data unravel the role of Pyk2 for modulating endothelial function, and remote cardioprotection in T2DM, highlighting the potential of Pyk2 as novel therapeutic target to re-establish impaired endothelium-dependent rIPC and thus optimize outcomes after AMI in diabetic conditions.

In particular, we showed (1) that NZO mice fully mimic key features of T2DM presented by insulin resistance, impaired glucose tolerance, hyperinsulinemia, hypertension and obesity; (2) this metabolic dysregulation was associated with severely impaired endothelium-dependent dilatory response to acetylchoine, carbachol, singular or repetitive limb ischmemic stimulation, impaired functional eNOS activity, reduced circulating NO bioactivity, increased arterial stiffness and blood pressure; (3) in non-diabetic humans and mice repetitive limb ischmia induced eNOS activation, flow-dependent dilation, and release of NO bioactivity into plasma transmitting signals essential for remote tissure protection in isolated hearts and aortic rings. In plasma of patients with T2DM and NZO mice, this protective signals through rIPC maneuvers were entirely lost. (4) In diabetic NZO mice, Pyk2 was highly activated, consecutively increasing the phosphorylation of eNOS on its inhibitory site Tyr656, which resulted in severe endothelial dysfunction and abolished endothelium-dependent remote protection (5) The inhibition of Pyk2 restored eNOS functional activity by dephosphorylation on Tyr656, resulting in the restoration of endothelial function and endothelium-dependent cardioprotection after rIPC in NZO mice to reduce IS after AMI. (6) Scavening of NO through CPTIO abolished this cardioprotective effect in Pyk2-inhibited NZO mice and healthy control animals, while addition of the NO donor nitrite re-established cardioprotection supporting the notion that endothelium-dependent transmission of NO bioactivity in plasma exerts remote tissue protection. (7) In addition to the vascular endothelium, neuronal signaling pathways are involved in rIPC, which is not affected by Pyk2-inhibition.

### Model characteristics, eNOS- and Pyk2-regulation

New Zealand Obese (NZO) mice have been established as a murine model of T2DM, reflecting the entire metabolic continuum ranging from obesity, an altered glucose levels, hyperinsulinemia, and insulin resistance to manifest T2DM ^28^. The insulin receptor downstream signaling, activates Akt and finally eNOS, known as the metabolic pathway of the insulin signal cascade. In particular, the stimulation of eNOS via adapter substrates leads to the activation of kinases, including PI 3-K/AKT, and consequently of eNOS phosphorylation on its stimulatory site Ser 1117 ^29^. As consequence, NO output is increased ^30, 31^. Therefore, one would expect the decrease of systemic hemodynamics and large arterial stiffness also in NZO mice. We here provide evidence, that NZO mice display isolated systolic hypertension, increase in total peripheral resistance and PWV values, featuring a dysbalance in large arterial compliance. This apparent paradox might be resolved by the notion that the balance between the antiatherogenic metabolic pathway and the mitogenic pathway is disturbed in insulin resistance. As consequence insulin induced eNOS activation is impaired in these conditions. Indeed, the sustained stimulation of the insulin receptor decreased cGMP levels and NO-dependent relaxation ^30, 32, 33^. Of note, the regulation of eNOS activity is complex, and several regulatory phosphorylation events are responsible for its activation/inhibition. Insulin was previously reported to induce the phosphorylation of eNOS on its inhibitory site Tyr657 mediated by Pyk2. This subsequently reduced eNOS capacity to generate NO, O_2_^-^ or citrulline ^14, 16, 34^. NZO mice present a reduced activation of eNOS on Ser1177 and an increased stimulation on Tyr656. Pyk2 activation was simultaneously increased, and eNOS functional capacity decreased. This illustrates how hyperinsulinemia and insulin resistance negatively influences eNOS. Proatherogenic factors associated with T2DM, such as oxidative stress ^35^, angiotensin II ^36^, and proteins of the MAPK family ^37^, can elicit the activation of Pyk2. As a consequence, the inhibition of Pyk2 should result in the restoration of eNOS function. Indeed, Pyk2-inhibition reduced eNOS phosphorylation on Tyr656 and restored the protein’s functionality, demonstrated by increased nitrite levels after rIPC-mediated activation. Our data imply a significant increase in Pyk2 activity, which inhibits eNOS functionality in T2DM conditions. Inhibition of Pyk2 restores eNOS function and its susceptibility to activation by shear stress.

### T2DM impairs endothelial function, remote cardioprotection and is restored by Pyk2-inhibition

The importance of the endothelium in contributing to remote protective signaling via increased NO bioavailability is widely discussed ^7, 38-41^. The rIPC maneuver is known to reduce IS ^7^ and fails to act protectively in clinical trials with T2DM patients with AMI ^39^. Therapeutic strategies to reverse T2DM induced alterations are lacking ^7^. T2DM and insulin resistance are associated with severe ED due to eNOS impairment ^42, 43^. Our in-vivo analyses of the endothelial function by FMD showed a poor response of ECs in NZO mice. Ex-vivo assessment of the endothelium-dependent relaxation validated these results. We and others have previously demonstrated that the pharmacological inhibition or gene knockdown of Pyk2 improved endothelial function and eNOS activity in vitro ^16, 25^. The data presented in this study are the first highlighting the crucial role of Pyk2 to regulate endothelial function, eNOS activity, and systemic hemodynamics in vivo, in a model of T2DM.

Multiple studies suggest that the endothelium releases several factors with a protective action on cardiomyocytes. Previously, we established that the endothelium partially contributes to rIPC-mediated remote cardioprotection via increasing circulating NO metabolites in the plasma ^11^. Indeed, our transfer experiments delineate that the plasma transports a transmitter signal for cardioprotection whose tissue-protective effects are abolished in diabetic conditions in mice and human plasma. Several mechanisms other than the endothelium are postulated to be involved in rIPC-mediated signaling for cardioprotection. Besides humoral and cellular factors, the neuronal pathway impacts remote cardioprotection ^44^ but may also be altered in diabetic conditions ^45^. Our data support the assumption that the neuronal pathways do not significantly impact IS, at least in this experimental model of T2DM.

In vivo results demonstrate per se an increase of IS, and abrogated rIPC-dependent decrease of IS in NZO mice, compared to Bl6 mice after myocardial I/R. Notably, the effect size of restoration of cardioprotection via rIPC after Pyk2-inhibition was considerable and achieved almost complete resolution of T2DM-associated enlargement of IS (see Figure 3). The regulation of eNOS-activation is blood-flow and time-course dependent ^25, 46^. The reduction of IS in Pyk2-inhibitor treated NZO mice after rIPC may also be modulated by the endothelium within the coronary circulation ^47^. IS were not reduced in Bl6 and NZO mice receiving Pyk2-inhibitor alone. These results support the notion that the endothelium remote the heart elicits protective effects. Our results obtained in the transfer of plasma from mice and humans subjected to rIPC bioassays validate that the interaction of Pyk2 and eNOS in the remote endothelium plays a decisive role in cardioprotection in T2DM. Taken together, our results favor the pivotal role of Pyk2-inhibition to limit IS via the reconstitution of endothelium-dependent remote cardioprotection in T2DM.

## Conclusion

Inhibition of Pyk2 holds the potential to act as a novel target to rescue endothelial-dysfunction in T2DM. This improves remote preconditioning and cardioprotection to limit IS after I/R. Further studies are essential to translate these findings into patients with T2DM and AMI.

## Abbreviations

AKT: protein kinase B
cAMP: cyclic adenosine monophosphate
cGMP: cyclic guanosine monophosphate
cPTIO: 2-4-carboxyphenyl-4,4,5,5-tetramethylimidazoline-1-oxyl-3-oxide
CV: cardiovascular
CVD: cardiovascular disease
DM: Diabetes Mellitus
EC: endothelial cell
ED: endothelial dysfunction
EMP: endothelial microparticle
eNOS: endothelial nitric oxide synthase
eNOS^−/−^ mice: endothelial nitric oxide synthase-knockout mice
ERK: extracellular signal-regulated kinases
FAK: focal adhesion kinase
FMD: flow-mediated dilation
GLUT4: glucose transporter type 4
I/R: ischemia/reperfusion
IL: interleukin
i.p.: intra peritoneal
IPC: ischemic preconditioning
IRS1: insulin-receptor substrate 1
LAD: left artery descending
MAPK: mitogen-activated protein kinase
MAPKp38: p38 mitogen-activated protein kinase
miRNA/miR: microRNA
mRNA: messenger RNA
N. fem. res.: Nervus femoralis resection
nNOS: neuronal nitric oxide synthase
NO: nitric oxide
NOS: nitric oxide synthase
NOX: nicotinamide adenine dinucleotide phosphate oxidase
PI3K: phosphatidylinositol 3-kinase
Pyk2: proline-rich tyrosine kinase 2
Pyk2-I: proline-rich tyrosine kinase 2-inhibition
rIPC: remote ischemic preconditioning
ROS: reactive oxygen species
Ser1177: serine 1177
sGC: soluble guanylate cyclase
STAT3: signal transducer and activator of transcription 3
STEMI: ST-segment elevation myocardial infarction
TNFα: tumor necrosis factor-alpha
T2DM: Type 2 Diabetes Mellitus

## Sources of Funding

This work was supported by the German Research Council (CRC 1116 to C.J., M.K., G.H.; and by a research grant of the Forschungskommission, Medical Faculty of the Heinrich Heine University Duesseldorf (to R.E).

## Disclosures

None

## Notes

### Competing Interest Statement

The authors have declared no competing interest.

