## Supplementary Information for "Inhibition of proline-rich-tyrosine kinase 2 restores cardioprotection by remote ischemic preconditioning in type 2 diabetes mellitus"

**Short title:** Restoration of remote cardioprotection in diabetes mellitus

***** These authors contributed equally

Corresponding author:

Ralf Erkens, MD and Malte Kelm, MD

Department of Cardiology, Pulmonology and Vascular Medicine

Medical Faculty Heinrich Heine University of Düsseldorf

Moorenstraße 5, 40225 Düsseldorf, Germany

### Expanded Methods

#### Materials

Unless otherwise specified, chemicals were purchased from Sigma-Aldrich Co. LLC. (Deisenhofen, Germany). Materials for western blotting were purchased from Life Technologies (Invitrogen, Darmstadt, Germany), Cell Signaling Technology (Massachusetts, EUA) DB Bioscience, and Abcam (Cambridge, UK).

#### Animals

All experiments were performed with 15-18 weeks-old male New Zealand Obese (NZO/HILtJ), C57BL/6J (BL6), and global eNOS knockout (eNOS$^{-/-}$) mice. All animal experiments were approved by the local animal ethics committee (LANUV, NRW, Germany) according to the European Convention for the Protection of Vertebrate Animals used for Experimental and other Scientific Purposes (Council of Europe Treaty Series No. 123). Animal care was carried out following the institutional guidelines, and all mice received ad libitum drinking water and a standard rodent diet.

Experimental planning and execution followed the ARRIVE recommendations [1]. NZO mice [2] were obtained from the German Diabetes Center (DDZ, Professor Dr. Al-Hasani, Germany). BL6 mice were purchased from the Jackson Laboratory (The Jackson Laboratory, Bar Harbor, ME, USA) or the Janvier Labs (Janvier Labs, Le Genest-Saint-Isle, France). Global eNOS$^{-/-}$ mice with C57BL/6J background were obtained from Professor Dr. Axel Gödecke, Heinrich Heine University of Düsseldorf, Düsseldorf, Germany [3]. All mice were allowed a minimum of seven-day acclimatization period to local vivarium conditions while they were housed on a typical 12/12h light/dark cycle at the central animal research facility of the Heinrich-Heine University (HHU), Düsseldorf, Germany

#### Pharmacological Pyk2 inhibition

Pharmacological Pyk2 inhibition was achieved by the inhibitor PF-431396 hydrate (Sigma-Aldrich, EUA). PF-431396 is a known inhibitor of PYK2 and FAK kinases at $IC_{50}$ = 11 and 1.5 nM, respectively. For experimental setup, it was diluted in a saline solution containing 2% DMSO and administrated at a concentration of 5 µg/g of body weight via intraperitoneal (i.p.) at least 15 minutes before the begging of each protocol performed as already published[4]. Control mice received vehicle (saline solution containing 2% DMSO).

**Patient recruitment for hybrid transfer experiments**

Six individuals with a medical history of cardiovascular diseases with type 2 diabetes mellitus (T2DM) and five participants without T2DM were included. Exclusion criteria were chronic kidney disease (GFR <30 ml/min), known liver disease, acute myocardial infarction, elevated CRP levels, or elevated WBC. Clinical studies were performed according to the ethics committee of the Heinrich-Heine University of Duesseldorf (ID2017034183 – 5903R) and the institutional guidelines. Written informed consent was obtained from all participants.

**Remote ischemic preconditioning (rIPC)**

The rIPC maneuver was performed by placing a vascular occluder (8 mm diameter, Harvard Apparatus, Harvard, Boston, MA, USA) over the murine lower proximal limb, as displayed schematically in Figure SI1. The cuff was inflated to 200 mmHg (Druckkalibriergerät KAL 84, Halstrup Walcher, Kirchzarten, Germany), leading to a total occlusion of the artery. . Following five minutes of occlusion, the cuff was deflated, and local tissue was allowed five minutes of reperfusion. This procedure was repeated four times.

In human participants,  ischemia was induced with a blood pressure measuring cuff (Welch Allyn GmbH, Hechingen, DE). The cuff was placed on the upper lower arm and inflated to 250 mmHg (minimum 50 mmHg over the systolic blood pressure). The pressure was kept constant for five minutes.Consequently, the cuff was deflated, permitting a five-minute reperfusion of the artery. This procedure was repeated four times.

**Blood glucose and plasma insulin measurement**

Plasma glucose levels was measured following 8-hours of fasting of the mice. After baseline measurement, the mice received glucose (1g/kgBW) i.p., as part of the glucose tolerance test (GTT). Consecutive measurements of the blood glucose (Contour XT; Bayer Health Care, Leverkusen, Germany) and plasma insulin (insulin mouse ultra-sensitive ELISA; DRG Instruments, Marburg, Germany) followed 15, 30, 60, 120, and 240 minutes after glucose administration by extracting small amounts of blood from the tail vein.

#### Assessment of cardiac function

In-vivo cardiac assessment was performed, as previously described by us [5], using a high-resolution ultrasound system (18-38 MHz; Vevo 2100/Vevo3100, Visual Sonics Inc., Toronto, Canada), and the analysis was carried out using the manufacturer's software. Echocardiography analyses were performed before and 24 hours after IRI. Left ventricular (LV) volume, stroke volume (SV), cardiac output (CO), and ejection fraction (EF) were calculated in B-mode by identification of maximal and minimal cross-sectional area using the manufacturer's software Vevo Lab 5.6.1 (Visual Sonics Inc., Toronto, Canada). All parameters (besides the EF) were normalized to body surface area. The body weight used for normalization was measured prior to echocardiographic analyses.

#### *In-Vivo* Assessment of FMD and PWV

Assessment of the endothelial function *in-vivo* by FMD was performed by a method previously described in our laboratory [5] with some modifications. After isoflurane anesthesia (1.5-2%) and achieving a stable cardiopulmonary state (heart rate: 400-500 bpm, breathing rate: ~100 breaths per minute, body temperature: 37°C), a 30-70 MHz linear array Microscan transducer (Vevo 2100/3100, Visual Sonics Inc., Toronto, Canada), set in a stereotactic holder, was positioned to visualize the arteria iliaca externa. A vascular cuff (8 mm diameter, Harvard Apparatus, Harvard, Boston, MA, USA) was placed on the same lower extremity of the mouse, distal of the transducer. After recording baseline images, the blood flow in the artery was ceased by inflating the occluder up to 250 mmHg (Druckkalibriergerät KAL 84, Halstrup Walcher, Kirchzarten, Germany), and the pressure was kept constant for five min. The occluder was then deflated, and several recordings of the vessel were performed during the consecutive vasodilation. Based on the recorded images during the occlusion and reperfusion, vessel diameter (%) changes at specific time-points were calculated as the ($diameter_{time\text{-}point}$ / $diameter_{baseline}$) x100. The difference in maximum dilation was calculated only from the vessel diameter values obtained after cuff deflations as

$$\Delta Max(\%) = \frac{diameter(max) - diameter(BL)}{diameter(BL)} \times 100.$$

Vascular stiffness was assessed through the pulse view velocity by a method previously described by us [5]. Briefly, the distance (ΔD) between the common carotid artery and the external iliac artery was divided by the time (ΔT) needed by the blood to travel between the two arteries, measured by B-mode-, M-mode- and PV-Doppler images.

**Aortic ring bioassay**

Aortae were isolated, cleaned from the adjacent fat tissue, and cut into 2-3 mm long rings. These were mounted on 300 µm wires in organ baths (Graz Glass Tissue Bath, 2 ml, Hugo Sachs Elektronik, March, DE) perfused with Krebs-Henseleit solution (K-H buffer: NaCl (118 mM), KCl (4.7 mM), $MgSO_4$ (0.8 mM 1,6), $NaHCO_3$ (25 mM), $KH_2PO_4$ (1.2 mM), glucose (10 mM), and $CaCl_2$ (2.5 mM 3,33)) equilibrated with a mixture of 95% $O_2$ and 5% $CO_2$ and calibrated for 60 minutes. The tension between the wires was measured by a transducer (F30 Force Transducer Type 372, Hugo Sachs Elektronik, March, DE), amplified by a PowerLab 8/30 (ADInstriments – Europe Head Office, Oxford, GB), and recorded with the LabChart (ADInstriments – Europe Head Office, Oxford, GB). The maximum contraction was determined by applying KCL (80 mM) to the baths. EC function was assessed as a response to cumulative doses of acetylcholine (Ach, 0.1 nM – 10 µM) or carbachol (0.1 nM – 10 µM) after precontraction with a bolus of the $\alpha_1$-adrenergic receptor agonist phenylephrine (Phe) corresponding to the half-maximal effective concentration ($EC_{50}$). The function of the VSMC was assessed in response to cumulative concentrations of sodium nitroprusside (SNP; 0.01 nM – 10 µM) and Phe (0.1 nM – 10 µM), respectively.

Transfer experiments with (non)conditioned murine and human plasma included some modifications of the standard organ bath protocol: blood was withdrawn via cardiac puncture in mice and venous puncture in humans before and after the rIPC maneuver. Plasma was separated after centrifugation in full heparinized tubes. Aortic rings were incubated with a plasma/KH solution and receiving continuously a mixture of 95% $O_2$ and 5% $CO_2$. In some experiments, endothelial function was tested in response to cumulative concentrations of carbachol (0.1 nM – 10 µM) to exclude any effect of Acetylcholine-receptor mediated stimulus.

### Determination of NO Metabolites in Plasma

Mice were anesthetized with isoflurane anesthesia (1.5-2%). Blood was taken by cardiac puncture and placed onto a tube containing 100 µL of NEM/EDTA/PBS (100 mmol/L N-ethylmaleimide and 5 µL of 0.5 mmol/L EDTA) solution. Subsequently, the blood solution was centrifuged at 3000G for two minutes at 4ºC. The plasma and red blood cells (RBCs) were separated, immediately frozen in liquid nitrogen, and stored at -80 until further analysis.

At the time of measurement, plasma samples were thawed on ice, and nitrite concentrations were quantified by a chemiluminescence detector (CLD 88 e, Eco Physics GmbH, Munich, Germany) as previously described [6]. Plasma nitrate concentrations were measured by high-performance liquid chromatography in ENO-30 (AMUZA INC, San Diego, USA), following the manufacturer's instructions. Acquired data were analyzed in eDAQ Powerchrome software (eDAQ, Warsaw, Poland) [6].

### Assessment of systemic hemodynamics

The hemodynamic parameters were assessed following the invasive close-chest method described previously [5]. The method uses a 1.4 F Millar pressure conductance catheter (SPR-839, Millar Instrument, Houston, TX, USA) positioned into the left ventricle (LV) through the right carotid artery. A Millar Box recorded the pressure and LV developed pressure, rate of pressure development ($dP/dt_{max}$), and rate of pressure decrease ($dP/dt_{min}$). Acquired data were analyzed by LabChart 7 (AD Instruments, Oxford, UK).

### Model of acute myocardial infarction and assessment of infarct sizes

The mice were randomly divided into one of the experimental groups shown in SI Figure 4. Analgesia was achieved with buprenorphine (0.1 mg/kg) 30 minutes before LAD ligation. Mice were intubated and kept on isoflurane anesthesia (induction 3% V/V and maintenance 2% V/V) via a rodent ventilator side port. Regardless of the protocol used, all groups were exposed to the exact duration of anesthesia. Heart rate was recorded by Electrocardiogram (ECG) recorder PowerLab 4.0 (AD Instruments, UK). The respiratory rate (about ~110-120 breaths/min) and the body temperature (37.5°C) were kept constant throughout the experiment. The thorax was opened, and myocardial ischemia was induced by the occlusion of the left descendent artery (Prolene suture 8.0) for 30 minutes. ST elevation on the electrocardiogram confirmed LAD occlusion success. After 30 minutes of ischemia, the occlusion was opened allowing LAD reperfusion for 24 hours [7]. Postoperative analgesia was achieved by buprenorphine (0.5 mg/kg BW) s.c. every 8 h until euthanasia.

Infarct sizes (IS) were determined by triphenyl tetrazolium chloride (TTC) staining as described [7]. After adequate anesthesia (ketamine 100 mg/kg BW and xylazine 10 mg/kg BW) and anticoagulation with heparin (250 IU i.p.), hearts were excised and perfused with 0.9% NaCl. LAD artery was occluded in the exact location, and 1% Evans blue dye was injected into the aortic root to delineate the area at risk (AAR) from the not-at-risk area. The hearts were frozen at −20°C for 60 min and serially sectioned into 1 mm slices; each slice was weighed and incubated in 1% TTC for 5 min at 37°C. Slices were photographed under a stereo microscope. AAR and non-ischemic areas were evaluated by computer-assisted planimetry (Diskus software, Hilgers Technisches Büro, Königswinter, Germany) by an observer blinded to sample identity. The size of the myocardial infarction is expressed as a percentage of the infarcted tissue area divided by the AAR.

**Assessment of murine cardiac function after treatment with (non)- conditioned human plasma**

Hybrid transfer experiments included perfusion of murine hearts (BL6 mice) with plasma from preconditioned healthy and diabetic individuals. These experiments followed a protocol with several modifications. Human plasma samples (4 ml), obtained at baseline levels and after rIPC-maneuver, were placed in 12 to14 kDa dialysis tubes (SpectraPor, Spectrum Europe B.V., Breda, the Netherlands) and dialyzed for 24 hours at 4°C against a 20-fold volume of modified Krebs-Henseleit buffer. Dialysates were then oxygenated and equilibrated to 37°C before use.

After adequate anesthesia with ketamine (100 mg/kg BW i.p.) and xylazine (10 mg/kg BW i.p.) and anticoagulation with heparin (250 IU i.p.), the hearts of BL6 mice were explanted, and the ascending aorta was cannulated and connected to a Langendorff apparatus (Hugo Sachs Electronics, March-Hugstetten, Germany). The equilibration phase started with retrograde perfusion with modified Krebs-Henseleit buffer (NaCl (118 mM), KCl (4.7 mM), $MgSO_4$ (0.8 mM), $NaHCO_3$ (25 mM), $KH_2PO_4$ (1.2 mM), glucose (5 mM), pyruvic acid (1.9 mM) and $CaCl_2$ (2.5 mM)) at a constant pressure of 100 mmHg.

Cardiac and coronary endothelial parameters were evaluated *ex-vivo* by Langendorff-apparatus as described [8]. A water-filled balloon connected to a pressure transducer was inserted through the mitral valve into the LV for the recording of isovolumetric LV developed pressure, its positive and negative first derivate (+dP/dt and -dP/dt) and the LV end-diastolic pressure (LVEDP). Consequently, hearts were subjected to 20 seconds of global zero flow ischemia to test the coronary reserve, followed by a five-minute recovery. Readouts of the hybrid transfer analyses were the derivates of the LVEDP (as measures for contractility), the LVEDP (for inotropy), and IS following zero-flow ischemia (for cardioprotection). Hearts were excluded from the analysis when they met one of the following exclusion criteria after the five-minute recovery phase: (I) coronary flow > 4 ml/min, (II) LV developed pressure < 50 mmHg, or (III) a coronary flow reserve revealed by transient ischemia < 70% of the baseline flow.

**Organ harvesting and molecular analysis**

Mice were anesthetized with 100 mg/kg ketamine (Ketanest®) and 10 mg/kg xylazine (Rompun®) via i.p. injection. Heparin (25.000 E.I. pyBraun, Germany) was administrated i.p. five minutes before blood taking. Blood samples were taken by cardiac puncture and immediately centrifuged at 800G for 10 minutes at 4ºC. Plasma samples were collected, frozen in liquid nitrogen, and stored in a -80 freezer until further analysis. After systemic perfusion with a cold phosphate-buffered solution (DPBS) without calcium and magnesium, pH 7.4 (Sigma Aldrich), organs were collected and immediately placed into tubes and frozen in liquid nitrogen and stored in a -80 freezer until further analysis.

**Western Blotting**

Heart samples were lysed with RIPA buffer (1% NP40, 0.1% SDS, 0.5% sodium deoxycholate in PBS) added to a mixture of protease and phosphatase inhibitors [9] and homogenized with a Tissue Ruptor (Qiagen, Hilden, Germany). Afterward, samples

were placed in an ultrasonic bath at 4°C for 10 minutes. After that, the samples were centrifuged at 13300G for 15 minutes at 4°C, and the supernatant was collected.
Total protein concentrations were determined by Bio-Rad DC Protein Assay (Bio-Rad Laboratories, USA). For immunoblot, 75-100μg of protein were loaded in NuPAGE™ 4-12% Bis-Tris, Pre-cast gels, following the manufacturer's instructions. After electrophoresis, proteins were transferred onto nitrocellulose membrane Hybond P 0,2 (Amersham Biosciences, Munich, Germany). The membranes were blocked for one hour with 2% Amersham ECL Prime Blocking Reagent in T-TBS (10mM Tris,100mM NaCl,0.1%Tween) followed by an overnight incubation at 4 °C with a mouse anti-eNOS (1:250; custom made from#624086) anti-eNOS/NOS typeIII antibody, stock:1mg/ml in PBS pH7.4, BD Bioscience, Erembodegem, Belgium), a rabbit anti-phospho-eNOS (Ser1177) (1:500; #9571, CellSignaling Technology, Cambridge, UK) and a mouse anti-GAPDH (1:5000 Abcam, Cambridge, UK). The primary antibodies were diluted in T-TBS 5% BSA 0,1% Tween. After washing, the membranes were incubated with HRP-conjugated anti-mouse or anti-rabbit (1:5,000; BD Biosciences). The bands were visualized by chemiluminescence with Western Blotting Detection Reagent (GE Healthcare Amersham™ ECL Select™) in Chemidoc imaging (Biorad Califórnia, EUA). Band intensity quantification was performed using ImageJ software.

Heart samples were pulverized in dry ice for immunoprecipitation, collected, and lysed in a lysis buffer. For eNOS precipitation, 2mg of protein lysate was incubated overnight with an anti-NOS III antibody (BD Bioscience 610297; 2 µg/mg protein lysate). After that, agarose immunoprecipitation was performed following the manufacturer's instructions. After elution, 25µl of the sample was loaded onto 8% SDS gels, and standard immunoblot protocol was performed. Membranes were incubated overnight with pY656 eNOS antibody (custom-made by Eurogentec, diluted 1:500 in Rotiblock). ECL detected the chemiluminescence signal. Phospho-Y656-eNOS/eNOS ratio was expressed as the fold change of BL6 mice.

**RNA isolation and rt-qPCR**

For RNA isolation, heart tissue was collected and placed into tubes containing 10x tissue amount of RNA later solution (Sigma Aldrich). The tubes were stored for 24 hours at room temperature and then stored in a -80 freezer until further analysis.

Heart samples were disrupted and homogenized with RTL lysis buffer (Qiagen, Germany) using a Retsch mixer (Mixer Mill MM 400) for 3 minutes. Afterward, tissue lysates were transferred to QIAsheredder cell-lysate homogenizer (Qiagen, Germany) and centrifuged at 14 000 G for 2 minutes at 4 degrees Celsius. RNA isolation was performed following the manufacturer's protocol (RNeasy® MiniKit-Qiagen). After sample elution, total RNA was quantified using a Nanodrop. Real-time qPCR was performed using TaqMan Array miRNA (ThermoFisher) to determine gene expression of NOS3 (Hs01574659_m1), PTK2B (Hs_00169444_m1), and the housekeeping gene RPLP0 (Mm00725448_s1). The relative expression of each gene was quantified by the $2^{-\Delta\Delta}$CT method.

**Statistical analysis**

The sample size was calculated prior by using G-Power V3.1. (Heinrich Heine University of Duesseldorf). Statistical analysis was done with GraphPad Prism 9 for Windows (Version 0.2.2(134)). Results are displayed as mean ± standard error of the mean (SEM) or mean ± standard deviation (SD). Statistical analyses were carried out either by unpaired or paired t-test for comparison of two groups or by 1-way and 2-way analysis of variance [10], respectively, as appropriate, when comparing more groups or treatments, followed by Bonferroni's or Sidak's post hoc tests. Where indicated, an unpaired Student's t-test was used to determine whether the two data groups differed significantly. The effect size of different treatments on the IS after IR injury was calculated as relative changes of IS to sham treatment within each strain. Normal distribution was tested by the D'Agostino-Pearson test. $p < 0.05$ was considered statistically significant.

**SI Figure 1: rIPC protocol for humans and mice**

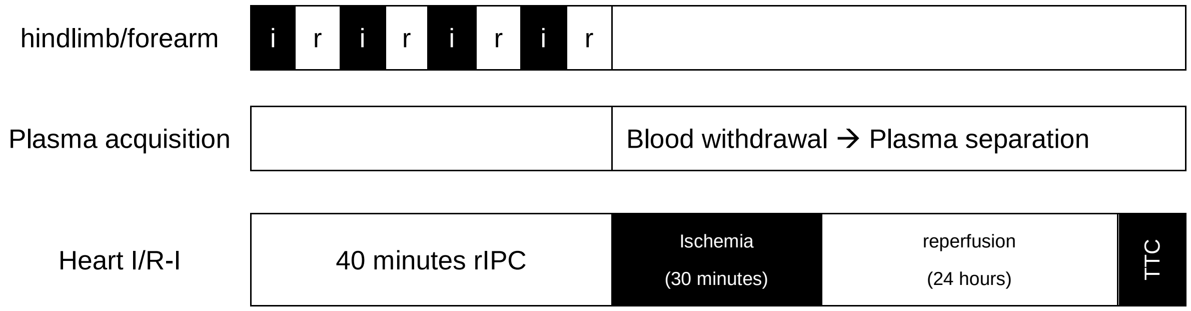


**SI Figure 1:** i = ischemia, r = reperfusion, I/R-I = ischemia-reperfusion injury, TTC = triphenyl tetrazolium chloride; rIPC = remote ischemic preconditioning.

**SI Figure 2: Functional eNOS activity is key for a proper endothelial-dependent vasodilation**

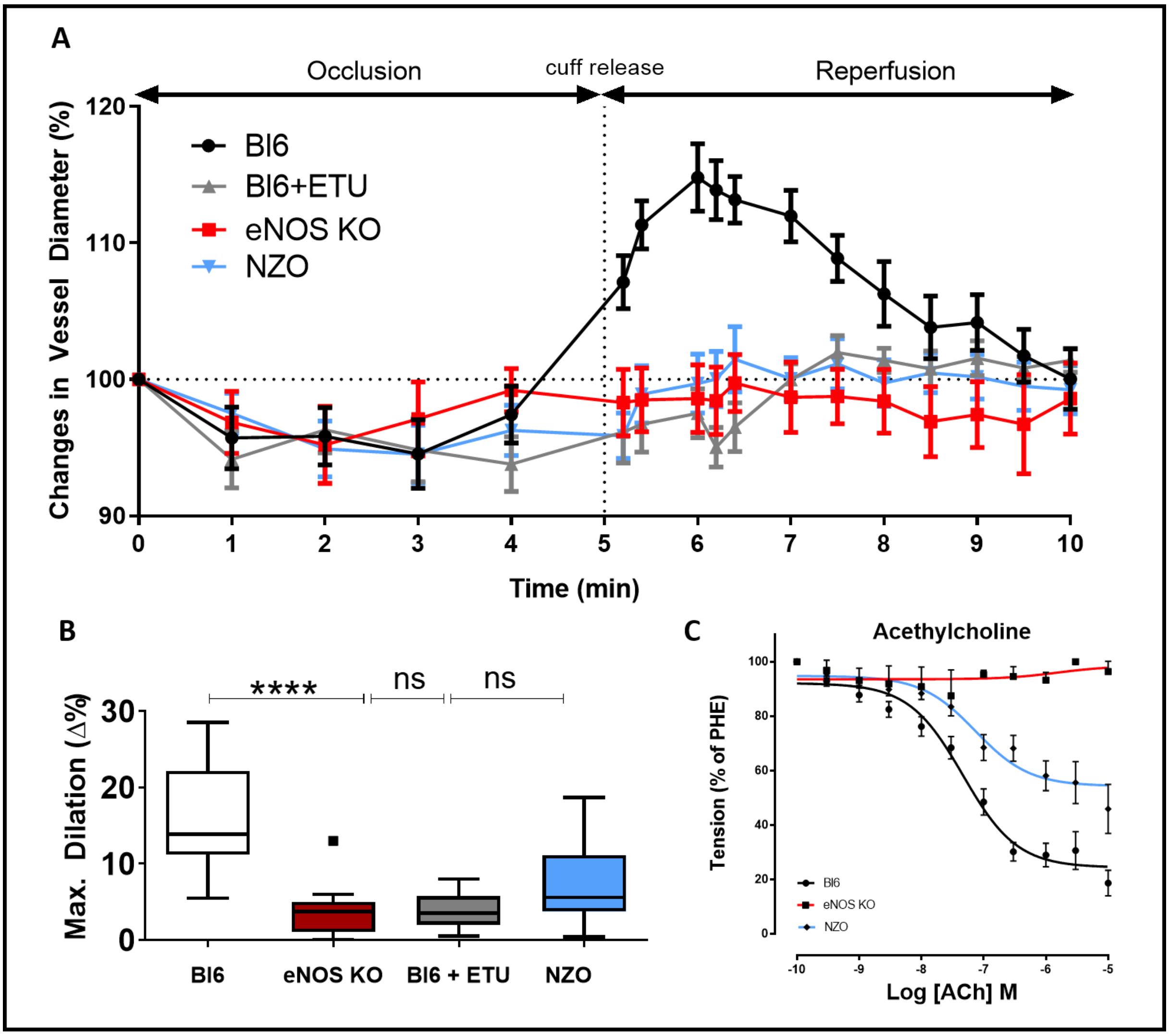


**SI Figure 2:** Endothelial function assessed: **(A)** *in-vivo* measurement of the flow-mediated dilation (FMD) in BL6 (black), BL6 treated mice with global eNOS-inhibitor ethylthiourea (ETU) (grey), eNOS-KO (red) and NZO mice (blue). (**B)** Maximal dilated diameter in FMD-response. (**C)** *ex-vivo* measurements of the endothelial function: eNOS$^{-/-}$ mice display a severely impaired endothelial relaxation response. *n* = 10–15; **** p < 0.0001.

**SI Figure 3: Supporting data to flow-mediated dilation and organ bath experiments**

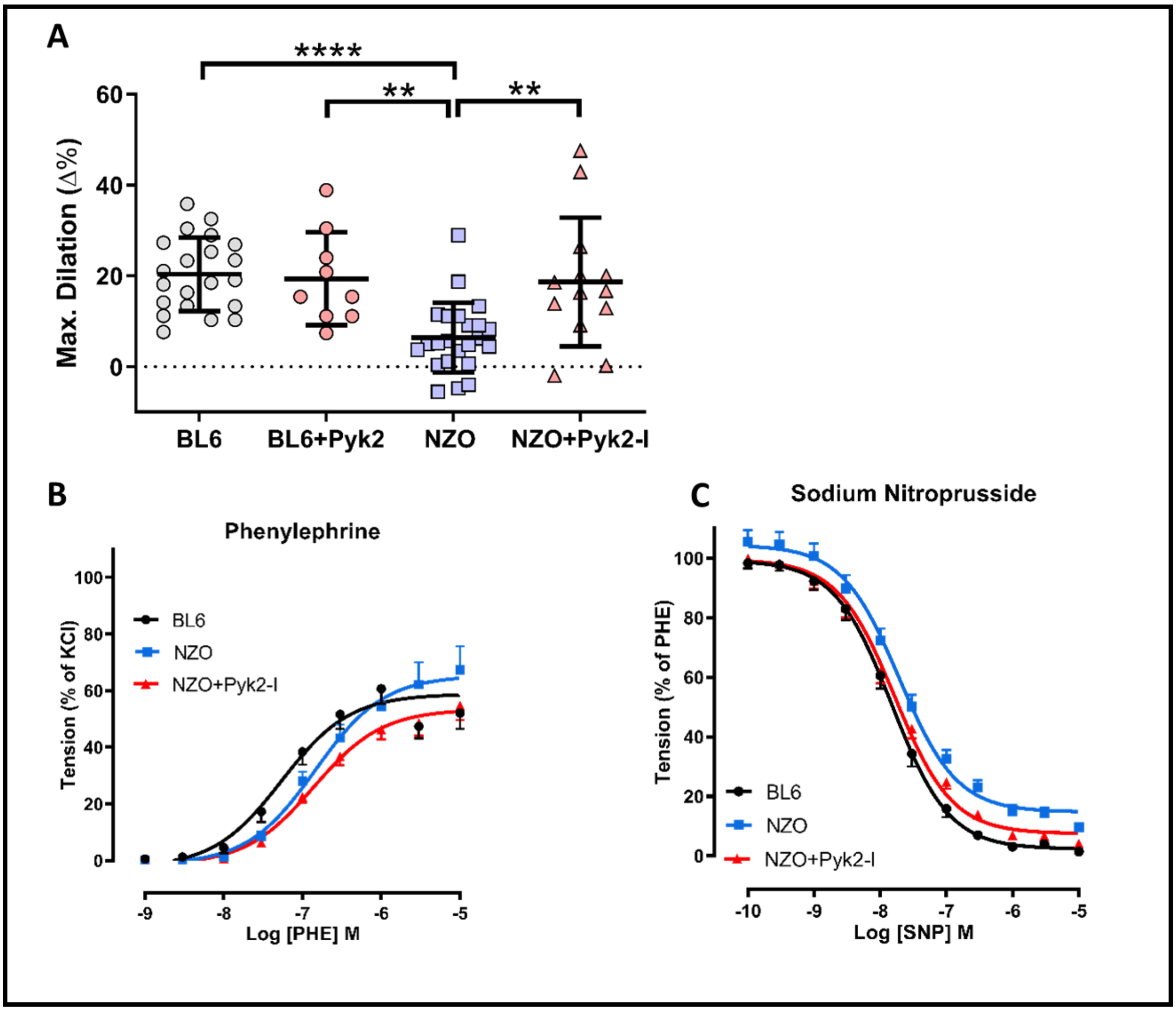


**SI Figure 3: (A)** Maximal dilated diameter in FMD-response: in BL6, BL6 treated mice with Pyk2-inhibitor NZO mice and NZO treated with Pyk2-inhibitor. **(B)** Phe-contraction and **(C)** SNP-relaxation curve corresponding to the organ bath experiments in Figure 3. $n$ = 10–15; ** $p$ < 0.01, **** $p$ < 0.0001.

**SI Figure 4: Schematic overview of protocols for ischemia and reperfusion injury protocols**

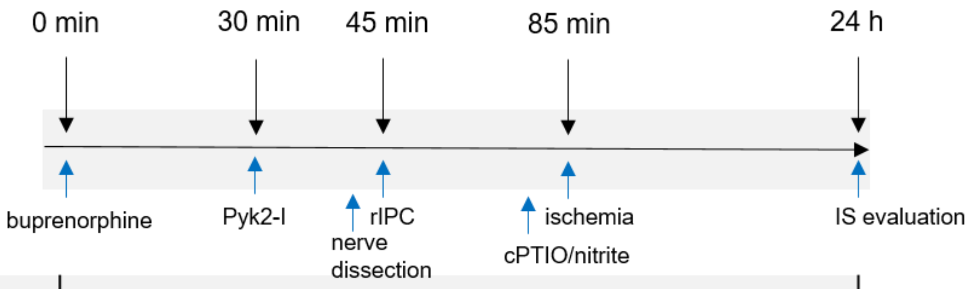


| | | | | |
|---|---|---|---|---|
| sham | | | | |
| rIPC | | | + | |
| Nitrite | | | | -/+ |
| rIPC + N. fem. res. | | + | + | |
| rIPC + cPTIO | | | + | +/- |
| Nitrite + cPTIO | | | | +/+ |
| Pyk2-I | + | | | |
| Pyk2-I + rIPC | + | | + | |
| Pyk2-I + Nitrite | + | | | -/+ |
| Pyk2-I + N. fem. res. + rIPC | + | + | + | |
| Pyk2-I + rIPC + cPTIO | + | | + | +/- |

**SI Figure 4:** Myocardial ischemia reperfusion injury consisted of 30 minutes of left-artery-descendend (LAD) ligation followed by 24h of reperfusion. In some experiments Pyk2-inhibitor was applied i.p. 15 minutes before performing the rIPC maneuver and 45 minutes before LAD ligation. Nervus femoralis resection (N. fem. res.) was performed five minutes prior to the rIPC maneuver and 25 minutes prior to LAD ligation. Exogenous nitrite supplementation or scavenging of endogenous NO bioactivity was achieved by administration of cPTIO or nitrite, which was performed i.p. 5 minutes before LAD ligation.

**SI Figure 5: The neuronal axis plays a secondary role in rIPC mediated cardioprotection both in healthy and diabetic mice**

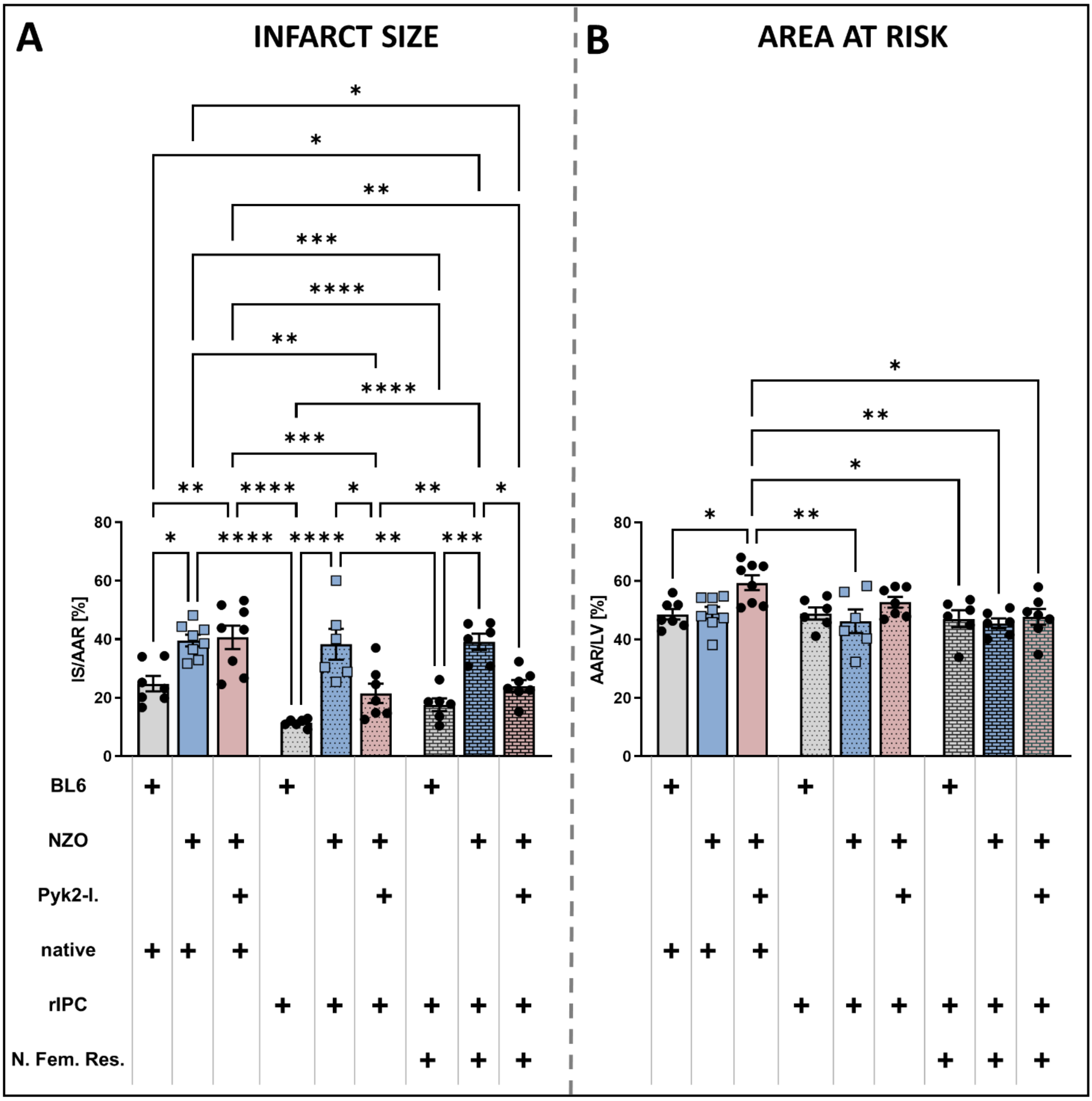


**SI Figure 5:** IS/AAR **(A)** and AAR/LV **(B)** of BL6 and NZO mice with and without Pyk2-inhibition (please refer to Figure SI 4 for detailed information on protocols). Nervus femoralis resection (N. fem. res.) was performed before performing the rIPC maneuver in each strain. In Bl6 mice rIPC conferred organ protection despite N.fem.res. In NZO mice, rIPC has no effect on the IS independently of nerve resection. After applying Pyk2-I. to NZO mice, the cardioprotective effects provided by rIPC were restored without changes by N. fem. Res. $n$ = 6–8; * $p < 0.05$; ** $p < 0.01$; *** $p < 0.001$; **** $p < 0.0001$.

**SI Figure 6: Areas at Risk per left ventricle corresponding to the infarct sizes shown in figures 4A and 5A.**

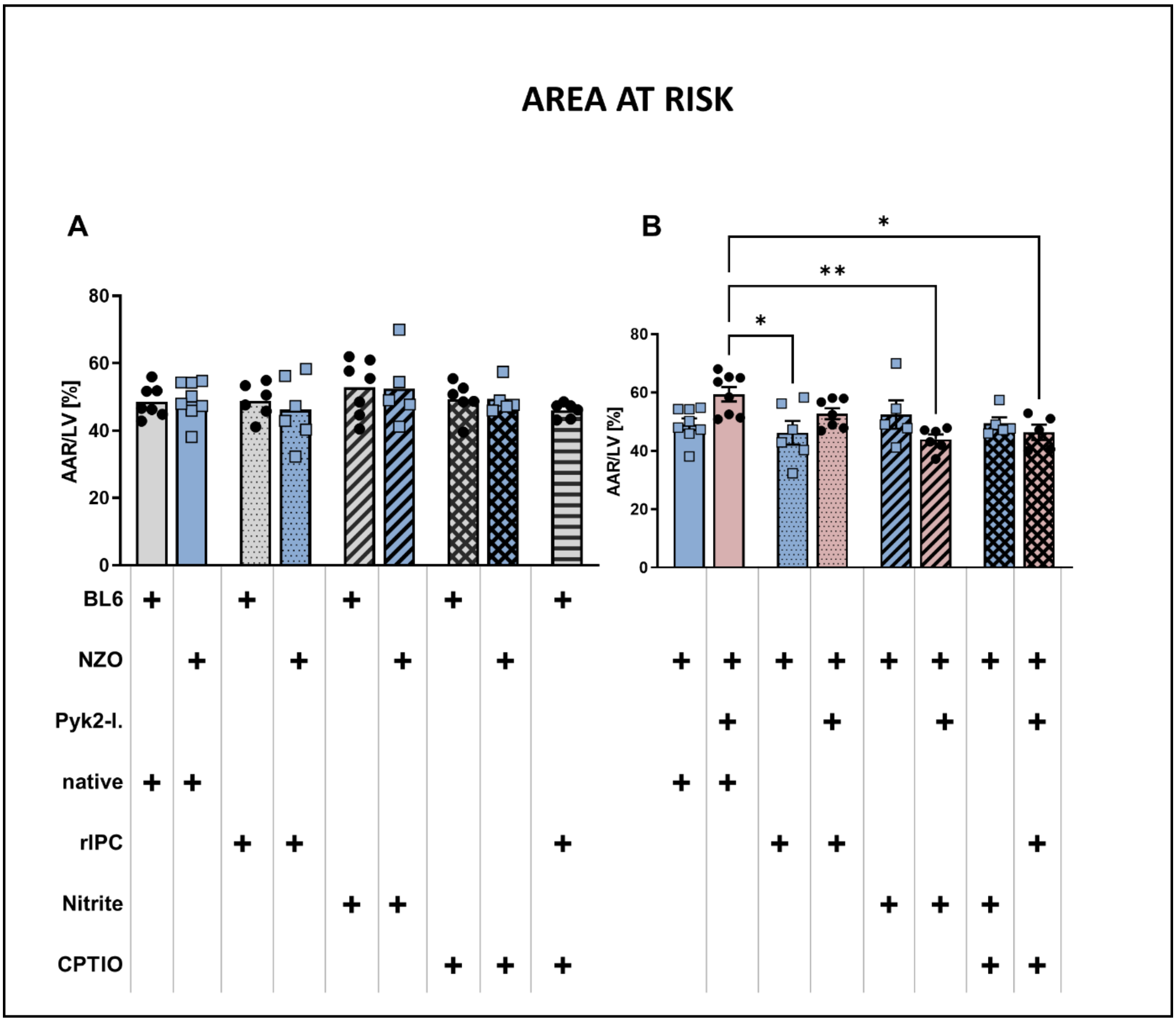


**SI Figure 5: (A)** AAR/LV comparison of BL6 vs. NZO mice shown in figure 4A. **(B)** AAR/LV comparison of NZO vs. NZO mice treated with the Pyk2-inhibitor shown in figure 5A. Distinct intervention per group is highlighted with a plus sign. Detailed information on protocols is shown in Figure SI 4. After I/R injury, AAR was not changed within most of the groups. $n$ = 6–8; * $p < 0.05$; ** $p < 0.01$.

**SI Figure 6: Supporting data to human (A) and murine (B) transfer organ bath experiments.**

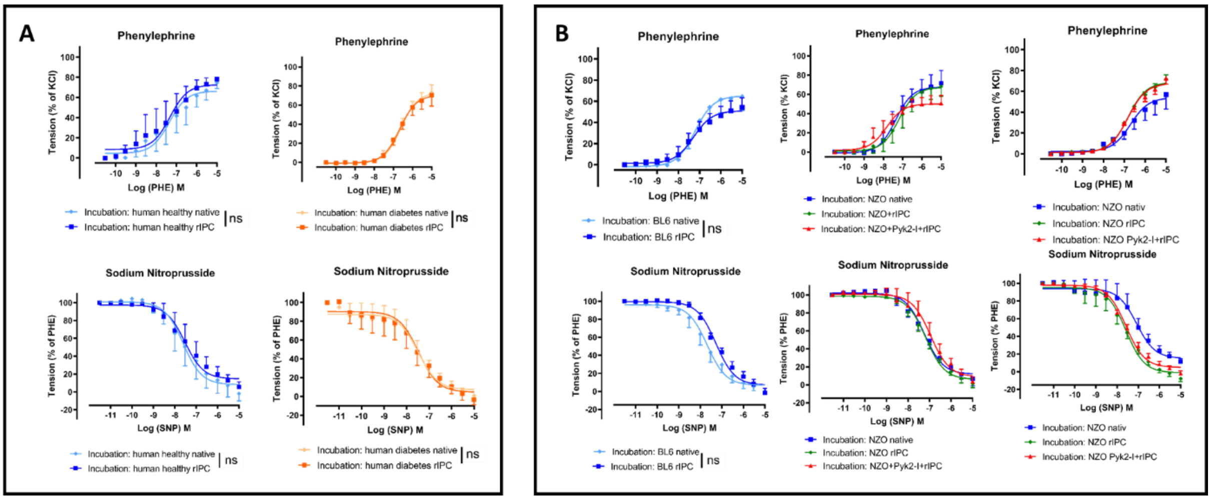


**SI Figure 6: (A)** Phe-contraction- and SNP-relaxation curve corresponding to the organ bath experiments in Figure 1, from transfer studies with human plasma. **(B)** Phe-contraction- and SNP-relaxation curve corresponding to the organ bath experiments in Figure 6, from transfer studies with murine plasma. $n = 6–8$; ** $p < 0.01$; **** $p < 0.0001$.

### SI Table 1: Patient characteristics

| Baseline characteristics | Patients with T2DM | Patients without T2DM | p-value |
|---|---|---|---|
| | n=6 | n=6 | |
| **Age, years** | 75.6 ± 6.2 | 79.8 ± 8.8 | 0.3765 |
| **Male, n[%]** | 5 [62.5] | 3 [60] | 0.9354 |
| **BMI, kg/m$^2$** | 28.6 ± 2 | 24.7 ± 1.75 | **0.0081** |
| Cardiovascular risk factors | | | |
| **Current smoking** | 1 [12.5] | 0 [0] | 0.5059 |
| **Hypertension** | 7 [87.5] | 3 [60] | 0.6236 |
| **Hypercholesterolemia** | 4 [50] | 0 [0] | 0.6236 |
| CVD parameters | | | |
| **Anemia** | 3 [37.5] | 2 [40] | 0.7114 |
| **Previous PCI** | 4 [50] | 0 [0] | 0.0979 |
| **Previous CABG** | 3 [37.5] | 0 [0] | 0.1877 |
| **SBP [mmHg]** | 125 ± 14 | 113.8 ± 8.2 | 0.2043 |
| **DBP [mmHg]** | 65.6 ± 8.8 | 65 ± 7.9 | 0.9317 |
| **Heart rate [bpm]** | 80 ± 14 | 67.5 ± 2.2 | 0.1456 |
| Laboratory parameters | | | |
| **HbA1c** | 8.9 ± 1.7 | 5.2 ± 0.07 | **0.001** |
| **Serum creatinine [mmol/l]** | 1.95 ± 0.94 | 1.2 ± 0.2 | 0.1384 |
| **Creatinine clearance [ml/min]** | 43 ± 29 | 60.4 ± 16 | 0.2873 |
| **Hb [g/dL]** | 11.6 ± 2.5 | 12.9 ± 0.91 | 0.3777 |
| **Sodium [mmol/L]** | 136 ± 5 | 138.8 ± 3.6 | 0.386 |
| **Potassium [mmol/L]** | 4.3 ±0.5 | 4.2 ± 0.3 | 0.7683 |
| **CK [U/L]** | 172 ± 95 | 111 ± 67 | 0.3467 |
| **NT-proBNP [pg/mL]** | 5714 ± 3869 | 1267 ± 1286 | 0.1668 |
| Concomitant medication | | | |
| **Insulin** | 7 [87.5] | 0 [0] | **0.0007** |
| **Biguanide** | 1 [12.5] | 0 [0] | 0.5059 |
| **DPP4 inhibitors** | 1 [12.5] | 0 [0] | 0.5059 |
| **Beta-blockers** | 7 [87.5] | 4 [80] | 0.5059 |
| **ACE inh. /AT-1 antag.** | 5 [62.5] | 3 [50] | 0.3596 |
| **Aspirin** | 5 [62.5] | 2 [40] | 0.7114 |
| **Statins** | 6 [75] | 2 [40] | 0.4332 |
| **Diuretics** | 8 100] | 1 [20] | **0.0012** |

**SI Table 2:** Baseline echocardiographic parameters of BL6, NZO and Pyk2-I treated NZO mice.

| | BL6 | NZO | | NZO+Pyk2-I | | |
|---|---|---|---|---|---|---|
| | mean ± SD | mean ± SD | p-value tested to BL6 | mean ± SD | p-value tested to BL6 | p-value tested to NZO |
| **Echocardiography** | | | | | | |
| n | 17 | 16 | | 7 | | |
| Heart rate (**HR**), bpm | 423.5 ± 42.3 | 427 ± 65 | 0.9821 | 414.5 ± 60 | 0.7861 | 0.7076 |
| Cardiac Output Index (**COI**), mL/min/cm² | 0.61 ± 0.15 | 0.6 ± 0.17 | 0.9734 | 0.57 ± 0.08 | 0.7661 | 0.8613 |
| Stroke Volume Index (**SVI**) µl/cm² | 1.46 ± 0.35 | 1.4 ± 0.3 | 0.8842 | 1.24 ± 0.09 | 0.2741 | 0.4692 |
| End-diastolic volume Index (**EDVI**), µL | 3.1 ± 0.65 | 2.88 ± 0.43 | 0.4356 | 2.68 ± 0.28 | 0.1838 | 0.6886 |
| End-systolic volume Index (**ESVS**), µL | 1.65 ± 0.46 | 1.47 ± 0.33 | 0.382 | 1.54 ± 0.17 | 0.8105 | 0.9088 |
| Ejection Fraction (**EF**), % | 47 ± 7.75 | 49 ± 8 | 0.7248 | 51.5 ± 2.2 | 0.3643 | 0.7222 |
| Interventricular Septum thickness end diastolic (**IVSd**), µm | 0.75 ± 0.12 | 0.92 ± 0.21 | **0.0095** | 0.86 ± 0.08 | 0.2444 | 0.7169 |
| Interventricular Septum thickness end systolic (**IVSs**), µm | 0.83 ± 0.11 | 1.15 ± 0.16 | **<0.0001** | 1.03 ± 0.15 | **0.0081** | 0.1481 |
| Left Ventricular Internal Dimension end diastolic (**LVIDd**), µm | 4.11 ± 0.28 | 4.55 ± 0.33 | **0.0004** | 4.46 ± 0.29 | **0.0397** | 0.7546 |
| Left Ventricular Internal Dimension end systolic (**LVIDs**), µm | 3.02 ± 0.37 | 3.17 ± 0.41 | 0.525 | 3.29 ± 0.49 | 0.3111 | 0.8051 |
| **Millar Catheter** | | | | | | |
| n | 8 | 9 | | 6 | | |
| Heart rate (**HR**), bpm | 504±51 | 486±55 | 0.7594 | 468.5±50 | 0.4396 | 0.8093 |
| Systolic Blood Pressure (**BPsyst**) - Aorta, mmHg | 91.75±9.4 | 112.1±12.6 | **0.0026** | 92±9.4 | 0.999 | **0.0057** |
| Diastolic Blood Pressure (**BPdia**) - Aorta, mmHg | 59.1±10.5 | 65.6±11.1 | 0.4259 | 47.8±8.9 | 0.1352 | **0.011** |
| Developed Pressure (**DP**), mmHg | 32.6±6.9 | 46.8±6.6 | **0.0003** | 43.8±2.8 | **0.007** | 0.6307 |
| End Systolic Pressure - LV, mmHg | 89.6±8.2 | 110.4±13 | **0.0011** | 88.8±6.8 | 0.9885 | **0.0017** |
| End Diastolic Pressure (**PED**) - LV, mmHg | 62.1±10.6 | 65.6±11.1 | 0.7806 | 48±92 | 0.0531 | **0.0124** |
| Maximum dP/dT (**$dP/dT_{Max}$**), mmHg/s | 3705±1208 | 4750±1174 | 0.1315 | 4733±509 | 0.1972 | 0.9995 |
| Minimum dP/dT (**$dP/dT_{Min}$**), mmHg/s | -1419±366 | -1920±433 | **0.0239** | -1791±157 | 0.1566 | 0.7772 |
